# Transport activity regulates mitochondrial bioenergetics and biogenesis in renal tubules

**DOI:** 10.1101/2024.02.04.578838

**Authors:** Chih-Jen Cheng, Jonathan M Nizar, Dao-Fu Dai, Chou-Long Huang

## Abstract

Renal tubules are featured with copious mitochondria and robust transport activity. Mutations in mitochondrial genes cause congenital renal tubulopathies, and changes in transport activity affect mitochondrial morphology, suggesting mitochondrial function and transport activity are tightly coupled. Current methods of using bulk kidney tissues or cultured cells to study mitochondrial bioenergetics are limited. Here, we optimized an extracellular flux analysis (EFA) to study mitochondrial respiration and energy metabolism using microdissected mouse renal tubule segments. EFA detects mitochondrial respiration and glycolysis by measuring oxygen consumption and extracellular acidification rates, respectively. We show that both measurements positively correlate with sample sizes of a few centimeter-length renal tubules. The thick ascending limbs (TALs) and distal convoluted tubules (DCTs) predominantly utilize glucose/pyruvate as energy substrates, whereas proximal tubules (PTs) are significantly much less so. Acute inhibition of TALs’ transport activity by ouabain treatment reduces basal and ATP-linked mitochondrial respiration. Chronic inhibition of transport activity by 2-week furosemide treatment or deletion of with-no-lysine kinase 4 (Wnk4) decreases maximal mitochondrial capacity. In addition, chronic inhibition downregulates mitochondrial DNA mass and mitochondrial length/density in TALs and DCTs. Conversely, gain-of-function Wnk4 mutation increases maximal mitochondrial capacity and mitochondrial length/density without increasing mitochondrial DNA mass. In conclusion, EFA is a sensitive and reliable method to investigate mitochondrial functions in isolated renal tubules. Transport activity tightly regulates mitochondrial bioenergetics and biogenesis to meet the energy demand in renal tubules. The system allows future investigation into whether and how mitochondria contribute to tubular remodeling adapted to changes in transport activity.

**Key points:**

- A positive correlation between salt reabsorption and oxygen consumption in mammalian kidneys hints at a potential interaction between transport activity and mitochondrial respiration in renal tubules.
- Renal tubules are heterogeneous in transport activity and mitochondrial metabolism, and traditional assays using bulk kidney tissues cannot provide segment-specific information.
- Here, we applied an extracellular flux analysis to investigate mitochondrial respiration and energy metabolism in isolated renal tubules. This assay is sensitive in detecting oxygen consumption and acid production in centimeter-length renal tubules and reliably recapitulates segment-specific metabolic features.
- Acute inhibition of transport activity reduces basal and ATP-linked mitochondrial respirations without changing maximal mitochondrial respiratory capacity. Chronic alterations of transport activity further adjust maximal mitochondrial respiratory capacity via regulating mitochondrial biogenesis or non-transcriptional mechanisms.
- Our findings support the concept that renal tubular cells finely adjust mitochondrial bioenergetics and biogenesis to match the new steady state of transport activity.

## Introduction

The kidney utilizes a wide variety of energy substrates to fuel robust transport activities across renal tubules to maintain fluid and electrolyte homeostasis (Wang et al., 2010). As one of the most energy-consuming organs, renal tubular cells have sizeable mitochondrial mass and oxygen consumption rate, especially the proximal tubule (PT), thick ascending limb (TAL), and distal convoluted tubule (DCT). Cumulative evidence has demonstrated that mitochondrial or nuclear gene mutations that aberrate mitochondrial function and metabolism are pathogenic in renal tubules, consequently debilitating transepithelial transports and cell survival (Kanako et al., 2022; Niaudet & Rötig, 2022; Viering et al., 2022; Viering et al., 2023).

Inhibition and stimulation of renal tubular transport activity by loop diuretics, thiazides, or vasopressin analogs decreased or increased, respectively, the mitochondrial mass and cell proliferation of TALs and DCTs (Bouby et al., 1985; Ellison et al., 1989; Kaissling et al., 1985; Loffing et al., 1995; Loffing et al., 1996). In mice, furosemide caused apoptosis and decreased the proliferation of TAL cells and, simultaneously, induced hyperplasia and proliferation of the downstream DCT cells. The correlation of histological and mitochondrial morphological changes with transport activity in TALs and DCTs leads to the hypothesis of “disuse atrophy” and “use hypertrophy” in TALs and DCTs, respectively. That is, transport activity, mitochondrial biogenesis/bioenergetics, and cell growth/proliferation are coupled. The role of mitochondrial function, such as mitochondrial respiration and metabolism, in these connections yet is not clearly defined, particularly with the resolution of a single tubular segment. Understanding mitochondrial respiration in renal tubules with different transport activities could provide further insights into the pathogenesis of renal tubulopathies, congenital or acquired, and the mechanisms of renal tubule remodeling (atrophy/hypertrophy, apoptosis/proliferation).

The major hurdle in studying renal mitochondrial metabolism is cell heterogeneity. The kidney has at least 16 types of renal epithelial cells featuring highly variable transport activities and metabolic features, with PT cells disproportionately outnumbering others (Balzer et al., 2022). Studies thus far using bulk kidney tissue cannot provide cell- or segment-specific information, especially for the distal part of the nephron. Assays using primary or immortalized renal cell lines hardly represent in vivo metabolism because the non-physiological culture environment inevitably changes the metabolic features of cultured cells, even for primary renal tubule cells (Dickman & Mandel, 1989; Johnson et al., 2017). Transport studies at individually isolated renal tubule level have revolutionized our understanding of kidney physiology and renal tubulopathies confined to a particular segment, such as Bartter’s or Gitelman’s syndromes. Our current knowledge of renal tubule metabolism came from early experiments using metabolic flux analysis of isolated renal tubules (Guder & Ross, 1984; Guder et al., 1986; Uchida & Endou, 1988). These studies revealed the distribution of metabolic enzymes along the nephron and how renal tubules metabolize diverse substrates for energy production or biosynthesis under different pathophysiological conditions.

Metabolic flux analysis using isotope-labeled carbon substrates requires an incubation time and is thus impossible for real-time measurements. Recent advances in “extracellular flux analysis (EFA)” allow us to measure oxygen consumption and proton production rates using sensor cartridges containing solid-state sensor probes to detect oxygen and proton levels in real time. These external metabolic fluxes can then be reductively interpolated into specific pathways when paired with metabolic substrate and inhibitor protocols. EFA has advantages in dynamic manipulations and measurements of mitochondrial respiration and glycolysis and has been popularly used to study energy metabolism in cancer cells and normal tissues, including kidney homogenates (McCrimmon et al., 2020). Whether EFA can be applied to isolated renal tubules and explore segment-specific mitochondrial respiration is unknown.

Here, we optimize and demonstrate that EFA can detect meaningful metabolic changes in centimeter-length renal tubules freshly isolated from mouse kidneys. We showed that isolated renal tubules are metabolically active and heavily rely on oxidative phosphorylation (OXPHOS) and, to a lesser extent, on glycolysis for ATP production. We then employed this method to show that acute inhibition of transport activity predominantly suppresses basal and ATP-linked mitochondrial respiration. Chronic alterations of transport activity further modify maximal mitochondrial respiratory capacity and mitochondrial mass. Our results establish a new approach for studying mitochondrial function in individually isolated renal tubules and show that mitochondrial metabolism may be a link between transport activity and tubular remodeling.

## Methods

### Ethical approval

All animal procedures were conducted in accordance with the University of Iowa’s guidelines for the care and use of laboratory animals. Institutional Animal Care and Use Committee (IACUC) at the University of Iowa has approved this study (approval number: 1082421-006). The investigators took all steps to minimize the pain and suffering of the animals throughout the study. The research complied with the ARRIVE guidelines 2.0 described in the Animal Ethics Checklist of *The Journal of Physiology* (Percie du Sert N et al., 2020).

### Animals

These experiments used 6∼8-week-old C57BL/6 wild-type, With-no-lysine 4 (Wnk4) knockout, and Wnk4^D561A/+^ knockin male mice weighing 20∼25 g. Mice were housed in cages made of polysulfone (Udel®, Thoren Caging Systems, Hazleton, PA) with dimensions of 19.56□×□30.91 × 13.34 cm. Cages are individually ventilated with humidity maintained between 30-70% and a temperature of 20-25°C under a 12:12-h light-dark cycle. Mice had ad libitum access to food (regular chow diet, Teklad Cat# 7913) and filtered water. For chronic furosemide treatment, age- and gender-match littermates were anesthetized with 2% isoflurane and underwent osmotic minipump (Alzet model 1002, Durect, CA, USA) implantation subcutaneously into the neck. The depth of anesthesia was determined using toe pinch. Briefly, mice were shaved at the back of the neck and scrubbed with betadine and alcohol. The pump was implanted subcutaneously along the left side of the back via an incision at the middle dorsal surface of the neck. The incision was closed with surgical staples. The dose of furosemide (30 mg/kg body weight/day, Sigma Aldrich, F4381) used in this study was the minimum effective dose used in previous studies (30-220 mg/kg) (Loffing et al., 1995; Na et al., 2003; van Angelen et al., 2012). The duration of furosemide treatment in this study (14 days) was longer than in previous studies (∼3-7 days) to achieve chronic suppression of the active transport in TALs. The mice’s body weight and urine osmolality were measured daily. Urine osmolality was measured using a micro-osmometer (Advanced Instruments, Model 210, Norwood, MA, USA). On Day 15, both kidneys and blood samples were harvested after the mice were sacrificed. All mice were humanely sacrificed by cervical dislocation, and they were not reused for other experiments.

### Microdissection of renal tubules

Renal tubules were microdissected as previously described (Cheng et al., 2012). Briefly, mouse kidneys were perfused with ice-cold modified Hank’s solution (composition (mM): 137 NaCl, 5 KCl, 0.8 MgSO4, 0.33 Na_2_HPO4, 0.44 KH_2_PO_4_, 1 MgCl_2_, 10 Tris(hydroxymethyl)amino methane hydrochloride, 0.25 CaCl_2_, 2 glutamine, and 2 L-lactate, pH 7.4, osmolality 295-300 mOsm/kg.H_2_O, bubbled with 95% O_2_/5% CO_2_) containing collagenase type I (1.5 mg/ml, Worthington, Lakewood, NJ, USA) and 1% bovine serum albumin. After perfusion, kidneys were harvested, sliced into thin sections, and incubated in a 15 ml tube with 10 ml pre-warmed collagenase-containing solution (same components of the perfusion solution) and then gently shaken on a titer plate shaker at 37°C for 15-20 minutes. The digestion time was strictly limited to less than 30 minutes to minimize cellular ATP depletion and cell death. After collagenase digestion, the individual nephron segments were dissected in ice-cold Hank’s solution pre-bubbled with 100% oxygen. The time for micro-dissecting PTs was no more than 60 minutes, while the time for TALs/DCTs was less than 150 minutes, based on the recommendations from a previous study (Uchida & Endou, 1988). The isolated renal tubules were transferred to Seahorse miniplates for EFA or 1% pre-warm agarose for transmission electron microscopy (TEM). In these experiments, all n are the number of mice and are as follows: (1) wild-type control (n = 84), (2) furosemide-treated (n = 6), (3) vehicle-treated (n = 6), (4) Wnk4 knockout (n = 6), (5) Wnk4^D561A/+^ knockin (n = 6).

### Extracellular flux analysis

Seahorse XFp and XF HS Mini analyzers were chosen for this study due to low sample size requirements compared to other XFe analyzers. The day before assays, the sensor cartridges were hydrated with calibrant and placed in a non-CO_2_ 37°C incubator overnight. On the assay day, Seahorse XFp miniplates were freshly coated with Cell-Tak solution (22.4 μg/ml), and the assay medium (Seahorse XF DMEM with 10 mM glucose, 1 mM pyruvate, and 2 mM glutamine or otherwise designated) and testing compounds were freshly prepared and pre-warmed in a 37°C bath following the manufacturer’s protocol. Isolated renal tubules were transferred to Cell-Tek coated miniplates, centrifuged at 300 x g for 2 minutes, and incubated with 100% O_2_ bubbled assay medium in a non-CO_2_ 37°C incubator for 15 minutes for PTs or 1 hour for TALs and DCTs prior to the assay. The incubation time was determined when the baseline measurements were stable without significant upward or downward drift in pre-testing. For acute inhibition of transport activity, isolated TALs were incubated with ouabain (500 μM, Sigma Aldrich, O3125) or vehicle (DMSO) in a non-CO_2_ 37°C incubator for 2 hours. Isolated renal tubules were evenly distributed in the center of the wells to ensure the accuracy of measurements.

Next, the overnight-hydrated sensor cartridge was loaded with pre-warmed testing compounds and transferred to the XF analyzer for calibration. The injection sequence and final concentration of testing compounds for each assay are listed below: Cell mito stress test: oligomycin A (inhibitor of mitochondrial ATPase, 3 μM for PTs/TALs, 6 μM for DCTs) in port A, carbonyl cyanide-p-trifluoromethoxyphenylhydrazone (FCCP, uncoupler of OXPHOS, 1 μM) in port B, and rotenone/antimycin A (Rot/AA, inhibitor of mitochondrial complex I and III, 0.5 μM) in port C; Real-time ATP rate assay: oligomycin in port A and Rot/AA in port B; Substrate oxidation stress test: UK-5099 (mitochondrial pyruvate carrier [MPC] inhibitor, 2 μM), BPTES (glutaminase inhibitor, 3 μM), or etomoxir (carnitine palmitoyl-transferase 1A [CPT1A] inhibitor, 4 μM) in port A, followed by a cell mito stress test with oligomycin-FCCP-Rot/AA in port B-D; Mito Fuel Flex test: glucose dependency: UK5099 in port A and BPTES/Etomoxir in port B, glucose capacity: BPTES/Etomoxir in port A and UK5099 in port B. The optimal concentrations of oligomycin and FCCP for each renal segment were titrated by separate pre-testing.

Before starting the assay, the miniplate was removed from the incubator and carefully washed with fresh pre-warmed assay media without detaching renal tubules. After assays, the results were analyzed by Wave 2.6 and Seahorse analytics software (Agilent Technologies, CA, USA). The following equations were used to calculate the ATP production rate: Glycolytic ATP production rate (pmol ATP/min) = glycolytic proton efflux rate (glycoPER, pmol H^+^/min) = total PER (pmol H^+^/min) –mitochondrial PER (mitoPER, pmol H^+^/min). (Total PER = ECAR [mpH/min] x buffer factor [2.5 mmol H^+^/L/pH] x Vol measurement chamber [2.28 μl] x volume scaling factor [Kvol: 1.19]; mitoPER = mitochondrial OCR (mitoOCR = OCR_basal_ – OCR_Rot/AA_) x CO_2_ contribution factor [CCF, 1.07 for PTs, 0.93 for TALs, and 0.94 for DCTs]). CCF, calculated as the ratios between mitoPER and mitoOCR, was determined by separate assays, in which isolated renal tubules were incubated in a glucose-free DMEM assay medium for 0.5 hours (PTs) or 1 hour (TALs & DCTs) to exhaust endogenous glycogen and glycolysis-dependent acidification. Mitochondrial ATP production rate (pmol ATP/min) = ATP-linked OCR (OCR_basal_ – OCR_oligo_, pmol O_2_/min) x 2 (pmol O/pmol O_2_) x the phosphate/oxygen ratio (P/O: ∼1.8 pmol ATP/pmol O in isolated mitochondria from rats’ renal cortex) (Schiffer et al., 2018; Edwards et al., 2020). The OCRs/ECARs were normalized by the total length (cm) of isolated tubules. In these experiments, all n are the number of mice and are as follows: (1) wild-type control (n = 78), (2) furosemide-treated (n = 6), (3) vehicle-treated (n = 6), (4) Wnk4 knockout (n = 6), (5) Wnk4^D561A/+^ knockin (n = 6).

### Quantitative PCR of microdissected renal tubules

To quantitate the relative content of mitochondrial DNA, total cellular DNA was extracted from the same amount of TALs (∼4cm) and DCTs (∼2cm) isolated from the experimental and control groups. Quantitative real-time PCR was conducted on C1000 Touch Thermo Cycler (Bio-Rad, CA, USA). Both mitochondrial (represented by mitochondrial NADH-ubiquinone oxidoreductase chain 1 [*Mt-nd1*] gene) and nuclear (represented by hexokinase 1 [*Hk1*] gene) DNAs were evaluated under the same experimental condition. The ratio of *Mt-nd1* to *Hk1* PCR products defines the relative mitochondrial DNA content. The total RNA of isolated renal tubules was immediately extracted using Quick-RNA™ MicroPrep kit (Zymo Research, Irvine, CA, USA). Reverse transcription was performed using Applied Biosystems™ High-Capacity cDNA Reverse Transcription Kit (Thermo Fisher Scientific, Waltham, MA, USA). We verified that no amplification was produced when reverse transcription was omitted from the sample. Relative mRNA abundance of genes of interest was standardized to the abundance of glyceraldehyde 3-phosphate dehydrogenase (*Gapdh*) mRNA and calculated by ΔΔCt value. Each specimen was assayed in triplicate. Sequences of primers for mitochondrial-to-nuclear DNA ratio and RT-PCR analysis in this study are *Aqp2* (forward: CTGGCTGTCAATGCTCTCCAC; reverse: TTGTCACTGCGGCGCTCATC), *Gapdh* (forward: CGTCCCGTAGACAAAATGGT; reverse: TCAATGAAGGGGTCGTTGAT), *Hk1* (forward: GAAAGGAGACCAACAGCAGAGC; reverse: TTCGTTCCTCCGAGATCCAAGG), *Mt-nd1* (forward: CTAGCAGAAACAAACCGGGC; reverse: CCGGCTGCGTATTCTACGTT), *Slc5a2* (forward: CAGACCTTCGTCATTCTTGCCG; reverse: GTGCTGGAGATGTTGCCAACAG), *Slc12a1* (forward: CCAGAGCGTTGTCTAAAGCA; reverse: TGGGCAGCTGTCATCACTTA), *Slc12a3* (forward: CTACCTTGCCATCTCAGCT; reverse: GGAGCATTCTGTGAAGTTCCAGC). In these experiments, all n are the number of mice and are as follows: (1) wild-type control (n = 32), (2) furosemide-treated (n = 12), (3) vehicle-treated (n = 12), (4) Wnk4 knockout (n = 7), (5) Wnk4^D561A/+^ knockin (n = 10).

### Electron microscope of isolated TALs and DCTs

Isolated TALs and DCTs were embedded into 1% agarose and fixed in 3% formaldehyde/glutaraldehyde (Electron Microscopy Sciences, Hatfield, PA) for 24 hours. The agarose blocks were then postfixed with 1% osmium tetroxide for 1 hour and rinsed in 0.1M sodium cacodylate buffer. After serial alcohol dehydration, the samples were embedded in Eponate 12 resin (Ted Pella, Redding, CA). After the polymerization of Epon, blocks were sectioned on an ultramicrotome, and ultrathin sections (70 nm) were poststained with uranyl acetate and lead citrate. Samples were examined by a Hitachi HT-7800 TEM (Tokyo, Japan). In these experiments, all n are the number of mice and are as follows: (1) wild-type control (n = 6), (2) furosemide-treated (n = 3), (3) vehicle-treated (n = 3), (4) Wnk4 knockout (n = 6), (5) Wnk4^D561A/+^ knockin (n = 6).

### Western blot analysis

The kidneys were harvested from mice immediately after sacrifice for Western blot (WB) analysis as previously described (Lin et al., 2021). Primary antibodies were applied overnight at 4°C, followed by secondary antibodies for 2 hours at room temperature. The primary antibodies used in this study include anti-Ncc antibody (Millipore Cat# AB3553, 1:5000 dilution), anti-Ncc phospho-Thr-58 antibody (1:2000 dilution), anti-Nkcc2 (Millipore Cat# AB2281, 1:2000 dilution), anti-Nkcc2 phospho-Thr-105 (Dundee Cat# S378C, 1:1000 dilution), anti-αENaC antibody (StressMarq Cat# SPC-403D, 1:2000 dilution), and anti-γENaC antibody (StressMarq Cat# SPC-405, 1:1000 dilution). The secondary antibodies used in this study include anti-rabbit (Jackson Lab Cat# 111-035-003, 1:4000 dilution) and anti-Sheep IgG antibody (Bethyl Cat# A130-101P, 1:10000 dilution). In these experiments, all n are the number of mice and are as follows: (1) wild-type control (n = 5), (2) furosemide-treated (n = 5).

### Statistical Analysis

A two-way ANOVA with mixed-effects analysis was applied to test for differences over time during 2-week furosemide or vehicle treatments in body weight and urine osmolality. A two-tailed Student paired t-test was performed to test for significant OCR response in oligomycin and FCCP titration tests. Simple linear regression was used to analyze the correlation between sample size (tubule length/Gapdh level) and readouts (OCR/ECAR). Nonlinear curve-fitting regression analysis was used to analyze the correlation between the calculated ATP production rate and sample sizes and the correlation between the percentage of glycolysis-driven ATP production and sample sizes. The comparisons between two groups were made using a two-tailed Student unpaired t-test. Mitochondrial morphology was quantified from the TEM images of TALs and DCTs taken from 3 different mice for each group (Lam et al., 2021). Mitochondrial volume density of TALs/DCTs was defined as the percentage of mitochondrial volume per renal tubular cell volume. The results from six tubules for each group were compared using a two-tailed Student unpaired t-test. To quantify mitochondrial length and cristae density, the long (major) axis and the inner mitochondrial membrane to outer mitochondrial membrane (IMM/OMM) ratio were measured in 50 mitochondria randomly picked from 6 tubules for each group. All TEM images were quantitated by using ImageJ (NIH). Data analysis and curve fitting were performed using Prism (v9.4.1) software (GraphPad Software, San Diego, CA, USA). Statistical significance was defined as p values less than 0.05. Data were presented as mean ± SD.

## Results

### Correlation of basal cellular respiration and acidification rates with the sample size of microdissected renal tubules in centimeter lengths

Specific renal tubular segments were manually dissected and validated by morphology and quantitative PCR of tubule-specific markers (*Slc5a2* for PTs, *Slc12a1* for TALs, *Slc12a3* for DCTs, and *Aqp2* for collecting ducts) (**Fig. 1**). To maintain cell survival and metabolism, we optimized the processes of sample preparation and pre-assay incubation, including a 30-minute restriction on collagenase treatment, oxygenated digestion/dissection solutions, ice-cold condition throughout tubule dissection, restriction on dissection time, and minimal incubation time before EFA assays (see methods). We first tested the sensitivity of EFA using different amounts of isolated PTs (0.3∼1.5 cm), TALs (0.7∼2.8 cm), or DCTs (0.2-0.9 cm).

**Figure 1.**
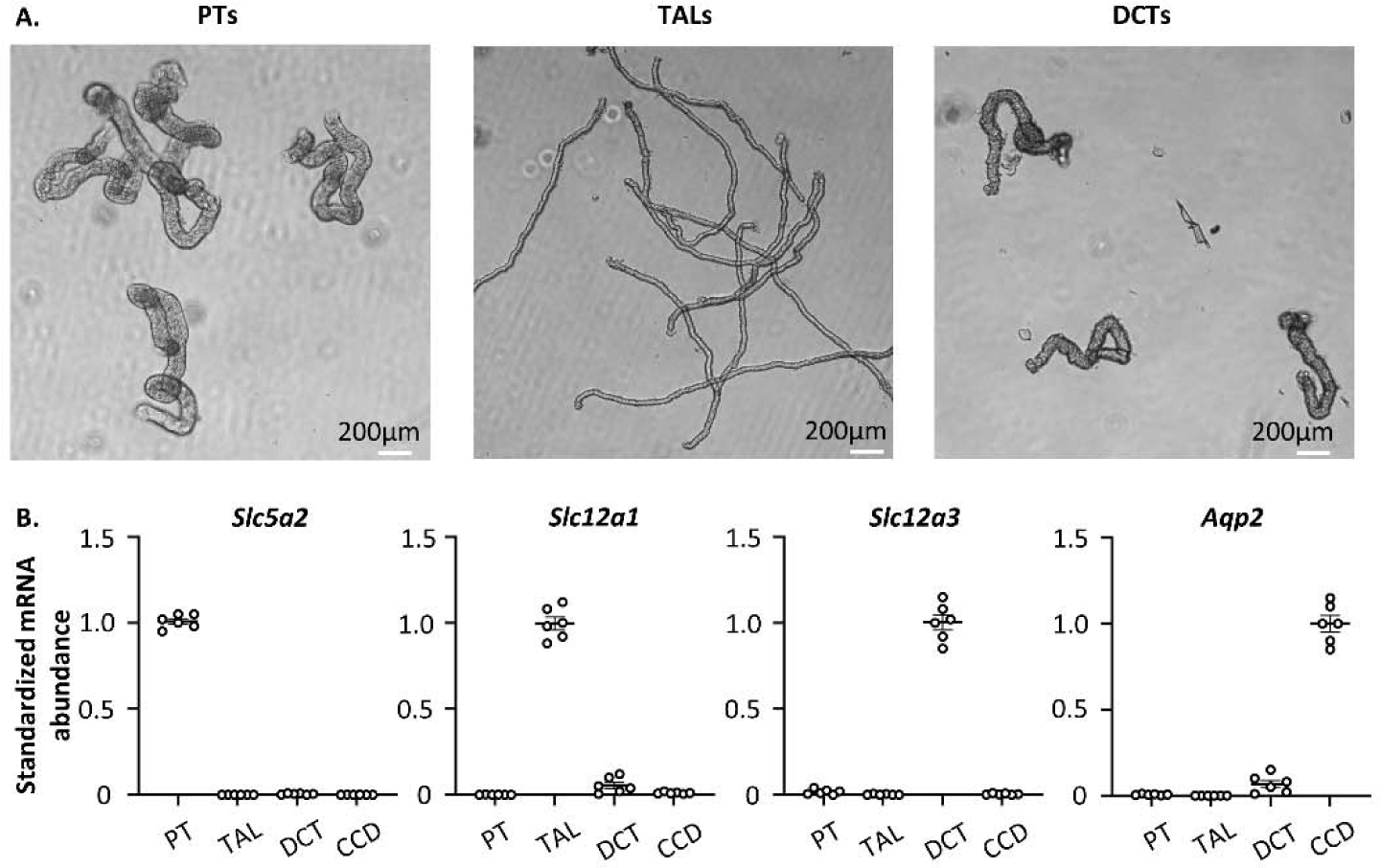
Authentication of isolated renal tubules. **(A)** The proximal tubules (PTs), thick ascending limbs (TALs), and distal convoluted tubules (DCTs) are recognized by their distinct gross appearance. PTs (S1 & S2) are large in diameter and convoluted. TALs are thin, straight, and parallel with S3 PTs and collecting ducts. The distal end of TALs is juxtaposed and attached to its parental glomerulus. DCTs are proximally connected to TALs, thicker in diameter than TALs, and distinctively convoluted. The distal ends of multiple DCTs merge into one collecting duct. **(B)** Segment-specific markers (*Slc5a2* for PTs, *Slc12a1* for TALs, *Slc12a3* for DCTs, and *Aqp2* for collecting ducts) were used to verify the purity of isolated renal tubules by quantitative PCR (n=6 for each segment).

The correlation analysis between basal OCRs and the total length of isolated TALs, DCTs, and PTs showed a positive linear association (**Fig. 2A-C**, left panel, n=3 tests for each segment). Alternatively, correlating OCRs with the relative abundance of the housekeeping *Gapdh* mRNA transcripts revealed similar results (**Fig. 2A-C**, right panels). We could not use protein concentration or cell numbers to estimate sample size due to the meager protein amount and cell superimposition in a 3D tubular structure. Basal ECARs were also positively associated with the total tubule length and Gapdh level of isolated TALs, DCTs, and PTs (**Fig. 2D-F**, n=3 tests for each segment). Compared to tubule length, Gapdh level did not yield a better correlation with OCRs/ECARs, especially in PTs. Therefore, we normalized OCR and ECAR results with tubule length in the following experiments. Next, we determined the optimal concentration of oligomycin and FCCP for each tubular segment. DCTs require a higher concentration (6 μM, p = 0.01) of oligomycin to suppress basal OCRs effectively than PTs and TALs (3 μM, p = 0.001 for PTs, p = 0.02 for TALs) (**Fig. 3A-C**, n=3 for each dose). FCCP 1 μM is optimal for uncoupling mitochondria and enhancing OCRs for all segments (p = 0.05 for PTs, p = 0.04 for TALs, p = 0.05 for DCTs) (**Fig. 3D-F**, n=3 for each dose).

**Figure 2.**
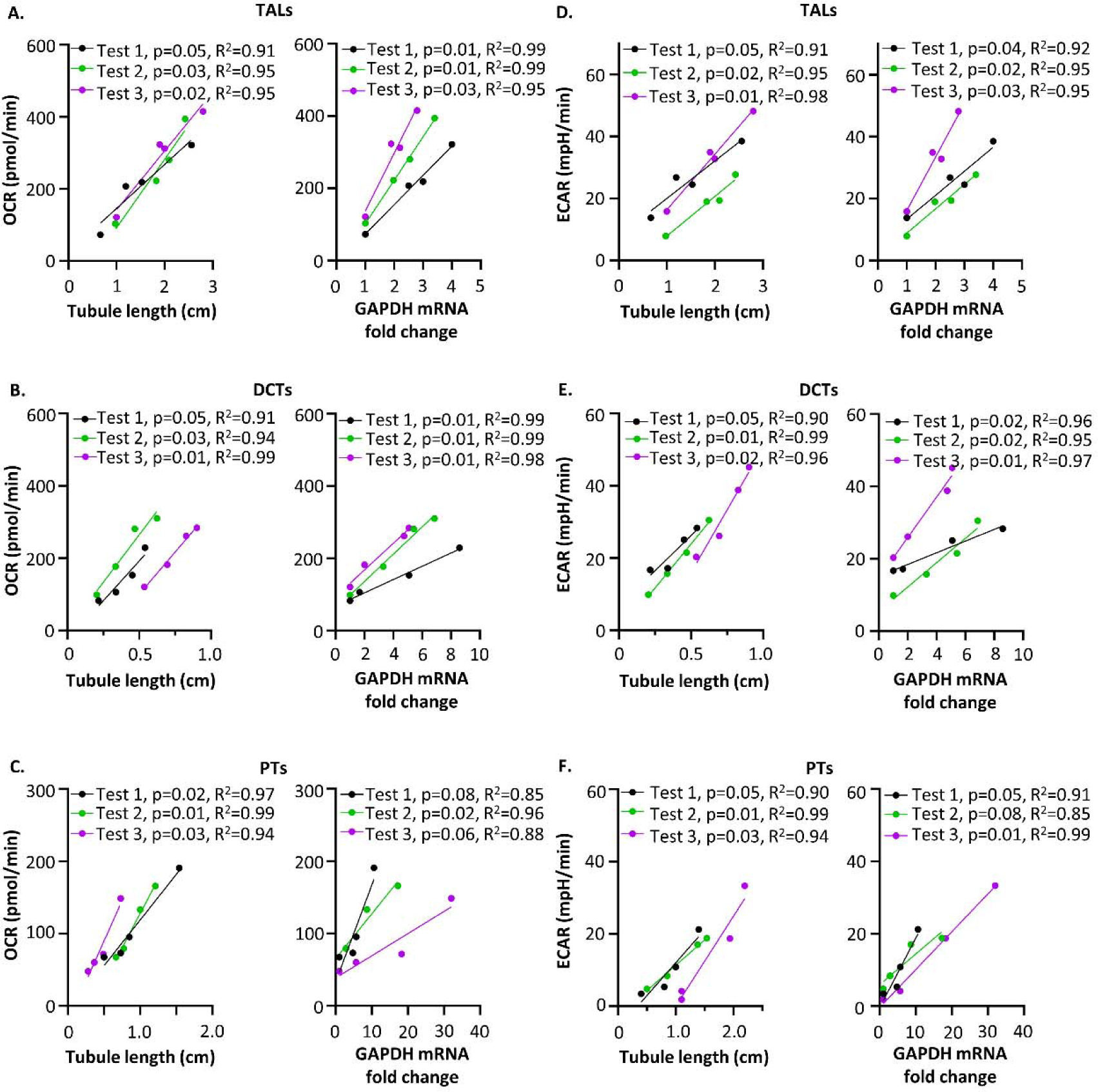
Basal oxygen consumption and proton production rates positively correlated with sample sizes of isolated renal tubules. Different amounts of isolated TALs, DCTs, and PTs were used for EFA assays to measure basal OCRs (**A-C**) and ECARs (**D-F**). Three separate EFA tests were conducted for each tubular segment (n = 3 for each segment). Each test included 4 renal tubule samples isolated from the same mouse kidney. After EFA assays, renal tubules were harvested for RNA extraction and quantitative PCR. The sample size was quantitated by the sum of lengths of isolated renal tubules using image J (left panel) or by the relative *Gapdh* mRNA level (fold-change versus the smallest sample) using quantitative PCR (right panel). The correlations between basal OCRs/ECARs and sample sizes were analyzed by simple linear regression. The p and R^2^ values for each test are shown in the figures. (DCT: distal convoluted tubule; ECAR: extracellular acidification rate; EFA: extracellular flux analysis; OCR: oxygen consumption rate; PT: proximal tubule; TAL: thick ascending limb)

**Figure 3.**
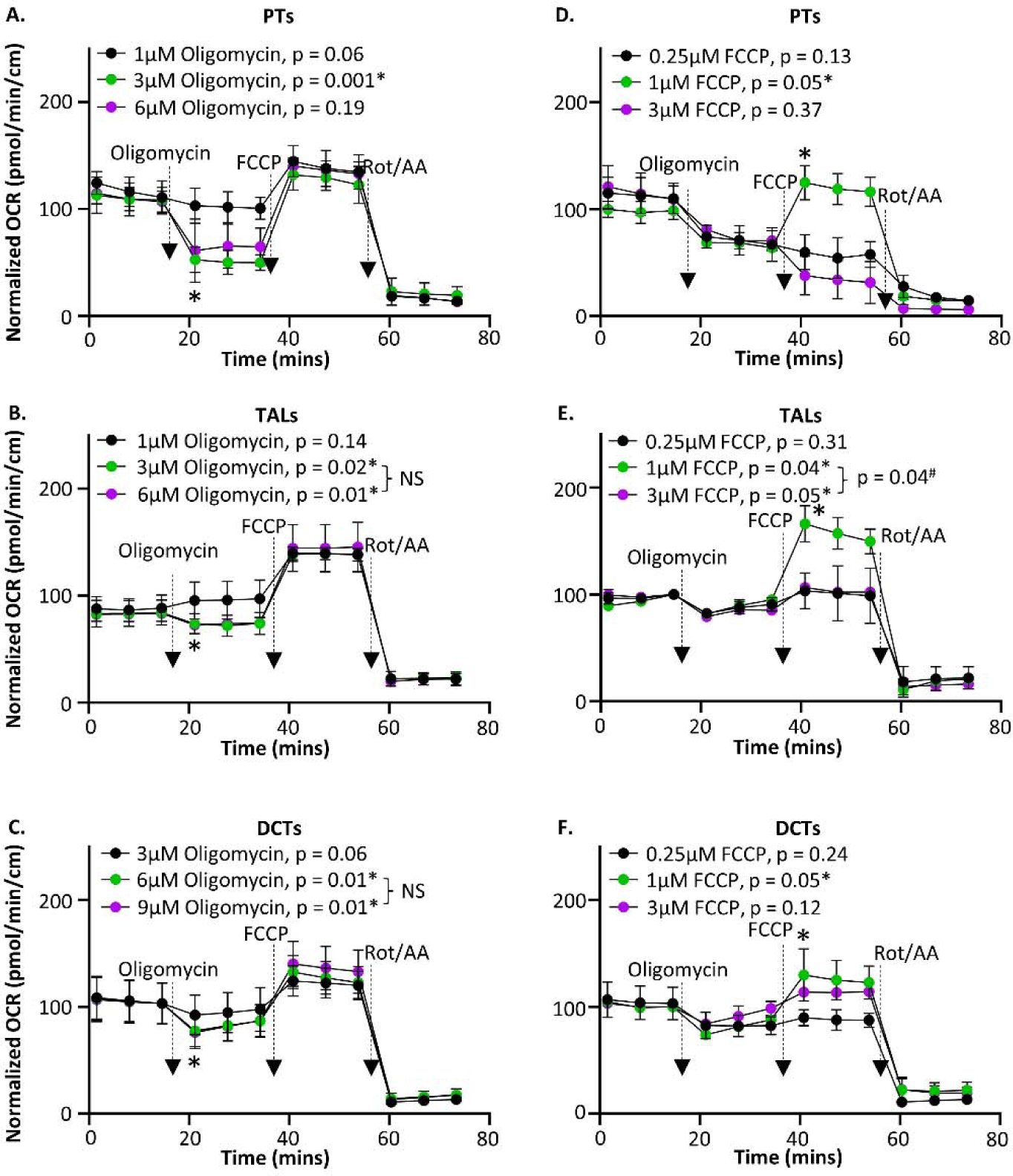
Optimization of oligomycin and FCCP concentrations for extracellular flux analysis using isolated renal tubules. Different doses of oligomycin or FCCP were used in Seahorse Cell Mito Stress Tests to determine the optimal oligomycin and FCCP concentration. **(A-C)** The optimal concentration of oligomycin for each tubular segment was determined when basal OCR was suppressed effectively by oligomycin (the 4^th^ vs. the 3^rd^ OCR) (n = 3 for each dose). *p < 0.05 (p values for each test are shown in figures) between the OCRs before and after oligomycin injections using a two-tailed Student paired-t test. NS: statistically not significant between two doses (TALs: 0.82, DCTs: 0.91) using a two-tailed Student unpaired t-test. **(D-F)** The optimal concentration of FCCP for each tubular segment was determined when OCRs were enhanced effectively (the 7^th^ vs. the 6^th^ OCR) by FCCP (n=3 for each dose). *p < 0.05 (p values for each test are shown in figures) between the OCRs before and after FCCP injections using a two-tailed Student paired-t test. ^#^p =0.04 between two doses using a two-tailed Student unpaired t-test. Data are presented as mean ± SD.

### Correlation of ATP production rate with the sample size of isolated renal tubules

The ATP production rate is physiologically relevant to cell bioenergetics and can be calculated from simultaneously measured OCR and ECAR values (Handel et al., 2019). OCRs measure mitochondrial OXPHOS, and ECARs measure acid efflux, thus reflecting lactic acid production and glycolysis. We performed the Seahorse real-time ATP rate assay to estimate the mitochondrial and glycolytic ATP production rates in isolated renal tubules. Injection of electron transport chain (ETC) complex inhibitors (oligomycin and Rot/AA) decreased OCRs and simultaneously increased ECARs in TALs and DCTs, indicating enhanced lactic acid formation and glycolysis when OXPHOS was inhibited in these segments (**Fig. 4A, B**, n=3 for each segment). In PTs, ECARs continued to decline when OCRs were suppressed, suggesting no significant compensatory glycolysis (**Fig. 4C**, n=3). The distinctive ECAR responses to OXPHOS inhibition are consistent with the fact that PTs have much lower glycolytic capacity than TALs and DCTs (Dickman & Mandel, 1989; Guder et al., 1986).

**Figure 4.**
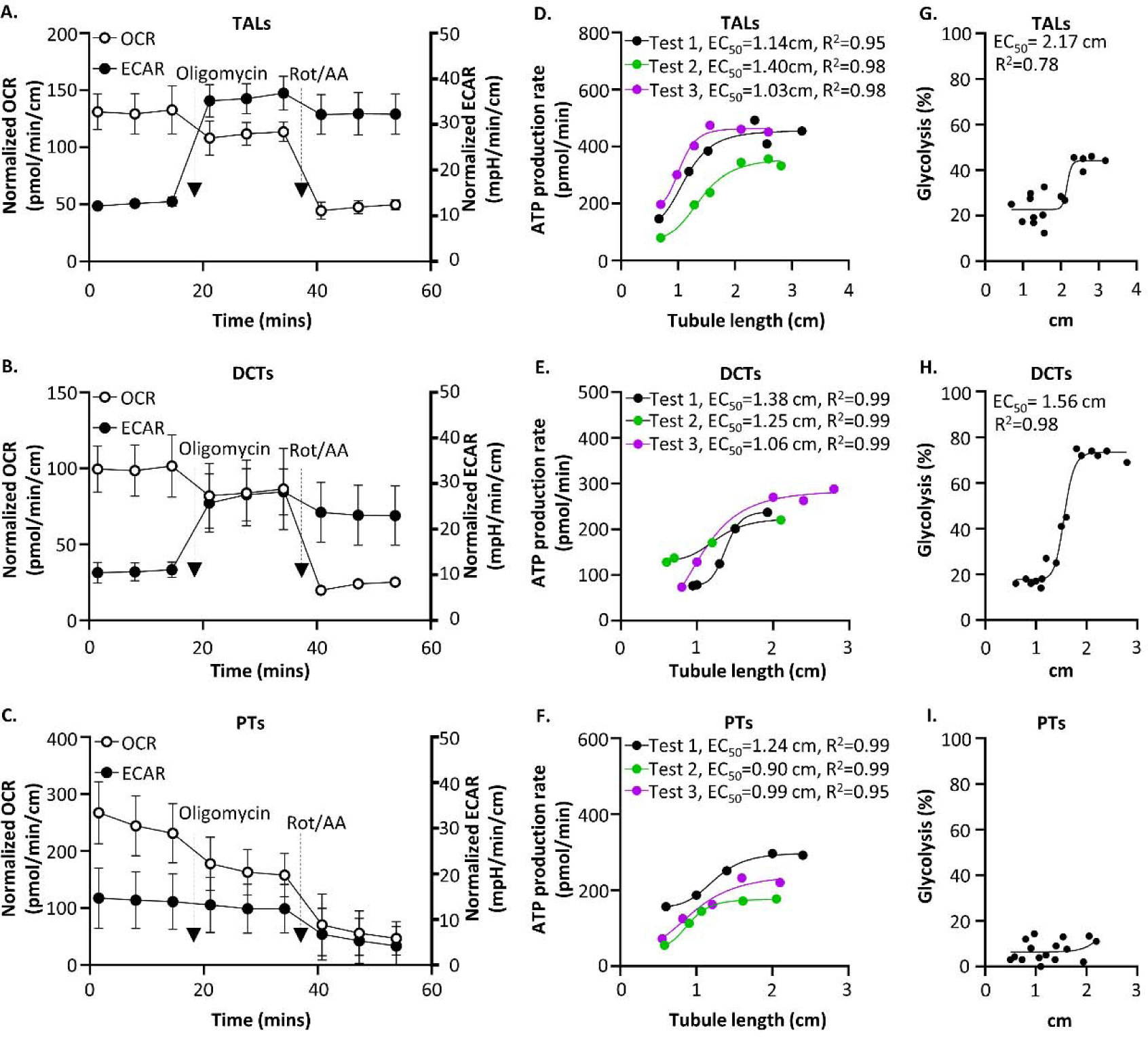
ATP production rates correlated with tubule length of isolated renal tubules before reaching a plateau. Real-time ATP rate tests were used to determine the relative contributions of mitochondrial OXPHOS and glycolysis to ATP production. For each test, 5-6 renal tubule samples with different lengths were isolated from the same mouse kidney. Three separate tests were performed. (**A-C**) A representative figure of simultaneous OCRs and ECARs measurements in real-time ATP rate tests using isolated TALs (**A**), DCTs (**B**), or PTs (**C**) (see supplemental Fig. 1 for the other two tests). OCRs and ECARs were normalized by tubule length. Data are presented as mean ± SD. (**D-F**) Calculated basal ATP production rates (see methods for equations) of isolated renal tubules from each test were nonlinearly correlated with tubule length for TALs (**D**), DCTs (**E**), and PTs (**F**) using nonlinear curve-fitting regression analysis ([Agonist] vs. response -- Variable slope (four parameters)). The EC_50_ (tubule length with OCR at 50% of the plateau) and R^2^ values for each test were shown. (**G-I**) The relative contribution of glycolytic ATP production to the total ATP production in TALs (**G**), DCTs (**H**), and PTs (**I**). Data from three tests were pooled for nonlinear curve-fitting regression analysis ([Agonist] vs. response -- Variable slope (four parameters)) (n=16 for each segment). The EC_50_ (tubule length with glycolysis percentage at 50% of the maximum [44% in TALs, 73% in DCTs]) and R^2^ values were shown in TALs and DCTs. Data from PTs did not reach a statistically meaningful result.

The calculated ATP production rates and the tubule length are positively correlated up to a certain threshold of tubular length (TALs: ∼2 cm, DCTs: ∼1.5 cm, PTs: ∼1.5 cm), and the ATP production rates reach a plateau (**Fig. 4D-F**, R^2^ ≥ 0.95, n=3 for each segment). Beyond that, the percentage of glycolytic ATP production over total ATP production significantly increases in TALs and DCTs (22.7 to 44.2% in TALs, R^2^ = 0.78; 17.5 to 73.5% in DCTs, R^2^ = 0.98) (**Fig. 4G, H**, n=16), indicating that oxygen supply relative to renal tubule abundance may be exceeded in these settings, therefore leading to increased glycolysis. Similar findings were observed in a study using brain tissue (Underwood et al., 2020). In contrast, overpopulated PTs do not increase the percentage of glycolytic ATP production, likely due to low glycolytic ability (**Fig. 4I**, R^2^ = 0.10, n=16). Based on these results, we recommend that the total tubule length for EFA assays should be less than 1.5 cm.

### Glucose is the primary energy substrate for TALs and DCTs

Many substrates can be oxidized for energy production by renal tubules. Several factors may affect substrate utilization in EFA assays, including sample preparation and substrate composition in assay media (Klein et al., 1981; Wittner et al., 1984; Mandel, 1985). Here, we investigated how TALs and DCTs utilize three major metabolic substrates, glucose, glutamine, and long-chain fatty acids (LCFA), to fuel mitochondrial respiration by sequentially injecting metabolic pathway inhibitors. UK-5099 is an inhibitor of mitochondrial pyruvate carrier (MPC), thus reflecting glucose dependence. BPTES is a glutaminase inhibitor, and etomoxir is an inhibitor of mitochondrial fatty acid carrier CPT1A.

UK-5099 reduced basal OCRs by 30% in TALs (**Fig. 5A**, glucose dependency test, n=3). Subsequent injection of BPTES and etomoxir thereafter did not further decrease OCRs, indicating the lack of compensatory utilization of glutamine and LCFAs when glucose oxidation is inhibited. Conversely, OCRs slightly increased when utilization of glutamine and LCFA was inhibited by BPTES and etomoxir but was inhibited by the subsequent MCP inhibitor UK-5099 (50% reduction in OCRs, **Fig. 5A**, glucose capacity test, n=3). Thus, mitochondrial respiration in TALs is ∼95% dependent on glucose oxidation and minimal dependence (< 5%) on glutamine or LCFAs oxidation (**Fig. 5B**). In DCTs, UK-5099 reduced basal OCRs by ∼20%, and the subsequent addition of BPTES and etomoxir caused an additional ∼4% OCR reduction (**Fig. 5C**, glucose dependency test, n=3). Conversely, BPTES and etomoxir together did not reduce basal OCRs, and the following UK-5099 significantly decreased OCRs (**Fig. 5C**, glucose capacity test, n=3). These findings suggest that DCTs mainly use glucose but may oxidize a small amount of glutamine and LCFAs when glucose oxidation is inhibited (∼16% flexibility). Besides, glucose can fully compensate for the deprivation of glutamine and LCFA oxidation (**Fig. 5D**).

**Figure 5.**
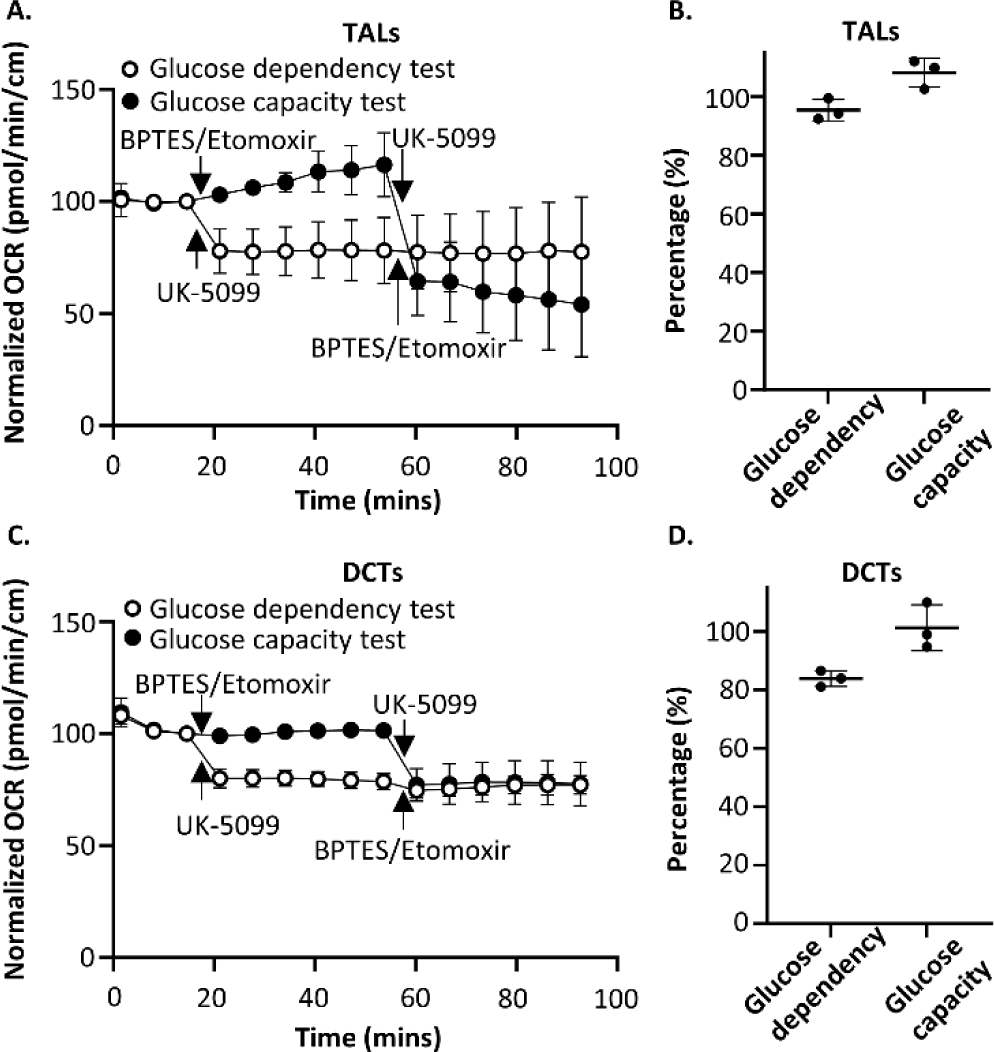
Mitochondrial fuel usage in the basal state of isolated TALs and DCTs. Seahorse XF Mito Fuel Flex Tests were used to determine the relative contributions of glucose, glutamine, and LCFAs oxidation to basal respiration in isolated TALs (**A, B**) and DCTs (**C, D**) by measuring OCRs in the absence and presence of pathway inhibitors. Glucose dependency and capacity tests were performed simultaneously with a reverse injection sequence of pathway inhibitors (see methods). For each test, six renal tubule samples were isolated from the same mouse kidney (n=3 for glucose dependency test, n = 3 for glucose capacity test). Three separate tests with similar results were performed for each tubular segment. (**A, C**) A representative OCR tracing of Mito Fuel Flex tests using isolated TALs (**A**) or DCTs (**C**) (see supplemental Fig. 2 for the other two tests). (**B, D**) The calculated glucose dependency and capacity of mitochondrial respiration in TALs (**B**) or DCTs (**D**). Results were summarized from three tests. Glucose dependency was calculated using the equation of [(mean OCR_basal (3rd point)_ – mean OCR_UK-5099 (4th point)_) / (mean OCR_basal (3rd point)_ – mean OCR_BPTES/Etomoxir (10th point)_)] x 100%, indicating the uncompensated part of mitochondrial respiration fueled by glucose oxidation. Glucose capacity was calculated using the equation of [1 – (mean OCR_basal (3rd point)_ – mean OCR_BPTES/Etomoxir (4th point)_) / (mean OCR_basal (3rd point)_ – mean OCR_UK-5099 (10th point)_)] x 100%, indicating the ability of renal tubules to oxidize glucose when the oxidation of glutamine and LCFAs are inhibited. All OCRs were normalized by tubule length and the 3^rd^ OCR (defined as 100 pmol/min/cm) of each sample. Data are presented as mean ± SD.

Critical substrate dependence is often revealed when cells are under high substrate demand, which happens when renal tubules have high active transport. It is difficult to stimulate transport activity in non-perfused renal tubules *ex vivo*. Therefore, we used mitochondrial uncoupler FCCP to enhance oxygen consumption and substrate oxidation to see whether TALs and DCTs oxidize glutamine and LCFAs to maintain mitochondrial proton gradient and ATP production. In TALs, UK-5099 suppressed basal and maximal OCRs by ∼35% (**Fig. 6A**, n=3). However, both BPTES and etomoxir had little effect on basal or maximal OCRs (**Fig. 6B, C**, n=3), suggesting high energy demand does not change TALs’ reliance on glucose under a condition of physiological glucose concentration (10 mM) (**Fig. 6D**). In DCTs, UK-5099 significantly, while BPTES and etomoxir slightly, reduced basal OCR. However, only UK-5099, not BPTES or etomoxir, significantly decreased the maximal OCRs (**Fig. 6E-G**, n=3), indicating that glucose is the primary energy substrate for mitochondrial respiration in basal and high energy demand states of DCTs (**Fig. 6H**).

**Figure 6.**
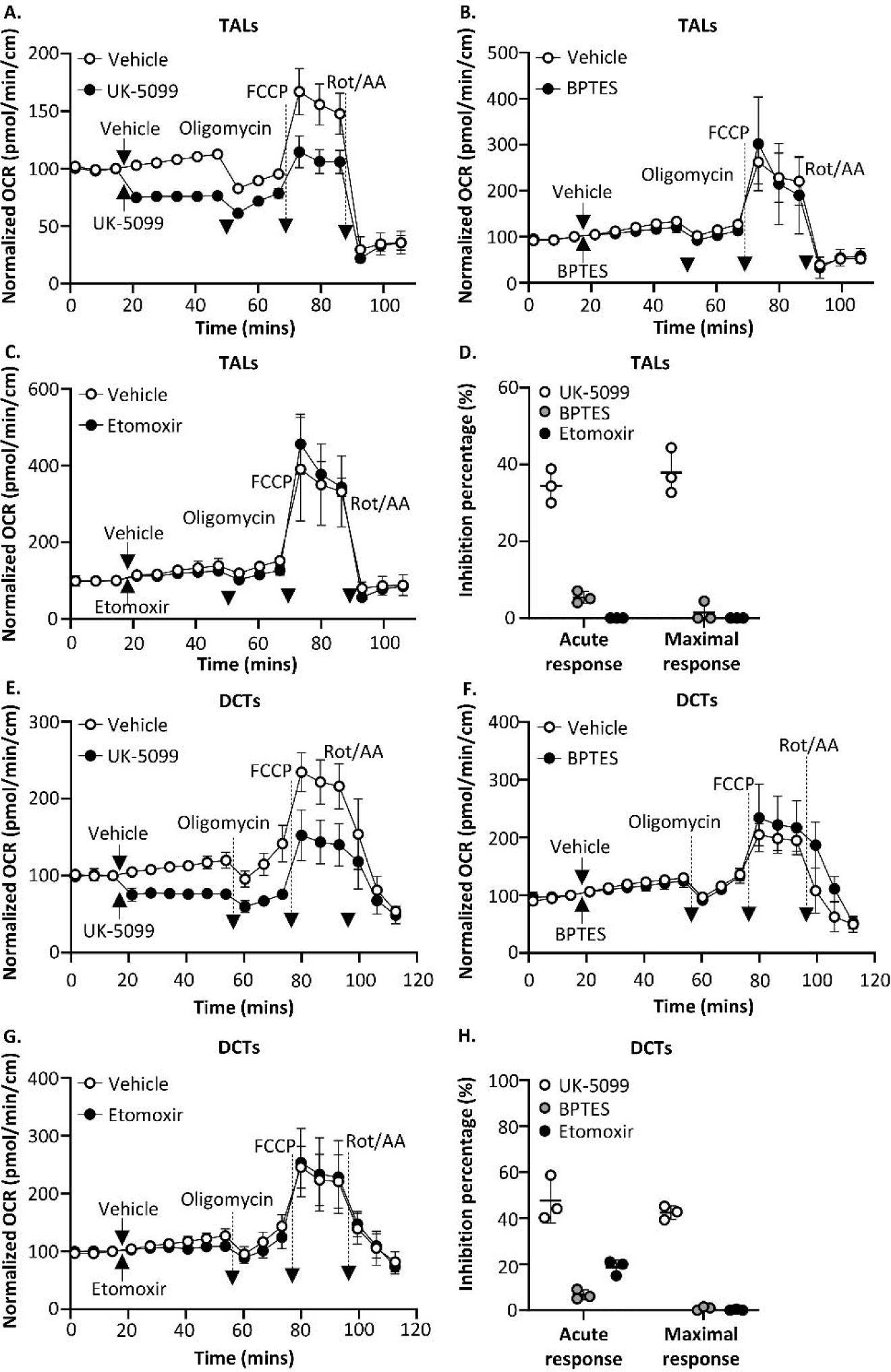
Mitochondrial fuel usage under high energy demand states of isolated TALs and DCTs. Seahorse XF Substrate Oxidation Stress Tests were conducted to investigate the oxidation of glucose, glutamine, and LCFAs in basal and high energy demand states in TALs (**A-D**) and DCTs (**E-H**). For each test, six renal tubule samples were isolated from the same mouse kidney (n=3 for the vehicle group, n = 3 for the metabolic inhibitor group). Three separate tests with similar results were performed for each tubular segment. (**A-C**) A representative OCR tracing of glucose (**A**), glutamine (**B**), or LCFAs (**C**) oxidation stress tests using isolated TALs (see supplemental Fig. 3A-C for the other two tests). (**D**) The relative contribution of glucose, glutamine, and LCFAs oxidation to basal (acute response) and maximal respirations (maximal response) in isolated TALs. Results were summarized from three tests. (**E-G**) A representative OCR tracing of glucose (**E**), glutamine (**F**), or LCFAs (**G**) oxidation stress tests using isolated DCTs. (see supplemental Fig. 3D-F for the other two tests). (**H**) The relative contribution of glucose, glutamine, and LCFAs oxidation to basal and maximal respirations in isolated DCTs. Results were summarized from three tests. Acute responses were calculated by [1 – (mean OCR_basal_ – mean trough OCR_URot/AA_ with pathway inhibitor) / (mean OCR_basal_ – mean trough OCR_URot/AA_ with vehicle)] x 100%, indicating the percentage of basal OCR being suppressed by pathway inhibitors. Maximal responses were calculated by [1 – (mean peak OCR_FCCP_ – mean trough OCR_Rot/AA_ with pathway inhibitor) / (mean peak OCR_FCCP_ – mean trough OCR_Rot/AA_ with vehicle)] x 100%, indicating the percentage of maximal OCR being suppressed by pathway inhibitors. The arrows mark the timepoint of pathway inhibitor or vehicle injection. The dashed arrows indicate the injections of ETC complex inhibitors and uncoupler FCCP. All OCRs were normalized by tubule length and the 3^rd^ OCR (defined as 100 pmol/min/cm) of each sample. Data are presented as mean ± SD.

### Transport activity regulates mitochondrial bioenergetics in TALs and DCTs

A positive linear correlation between urinary sodium reabsorption and oxygen consumption rate exists in mammalian kidneys, suggesting potential coupling between transport activity and mitochondrial respiration in renal tubules (Deetgen & Kramer, 1961; Mandel & Balaban, 1981). Having established EFA as a valid assay for mitochondrial respiration, here we acutely or chronically modulated tubular transport activity and measured the mitochondrial respiration in isolated TALs or DCTs.

Furosemide was given to mice via minipump for 2 weeks to suppress TAL transport activity chronically, reflected by the reduced mice body weight and urine osmolality from the first day of furosemide treatment (p values: body weight = 0.02, urine osmolality < 0.0001)(**Fig. 7A, B**). Western blot analysis of whole kidney lysates demonstrated enhanced protein expression of Ncc (total and phosphorylated forms, p values: Ncc = 0.01, p-Ncc = 0.02) and ENaC subunits (p values: cleaved ENaC-L = 0.04, ENaC-α = 0.05), indicating compensatory stimulation of active transport in DCTs and collecting ducts (**Fig. 7C**).

**Figure 7.**
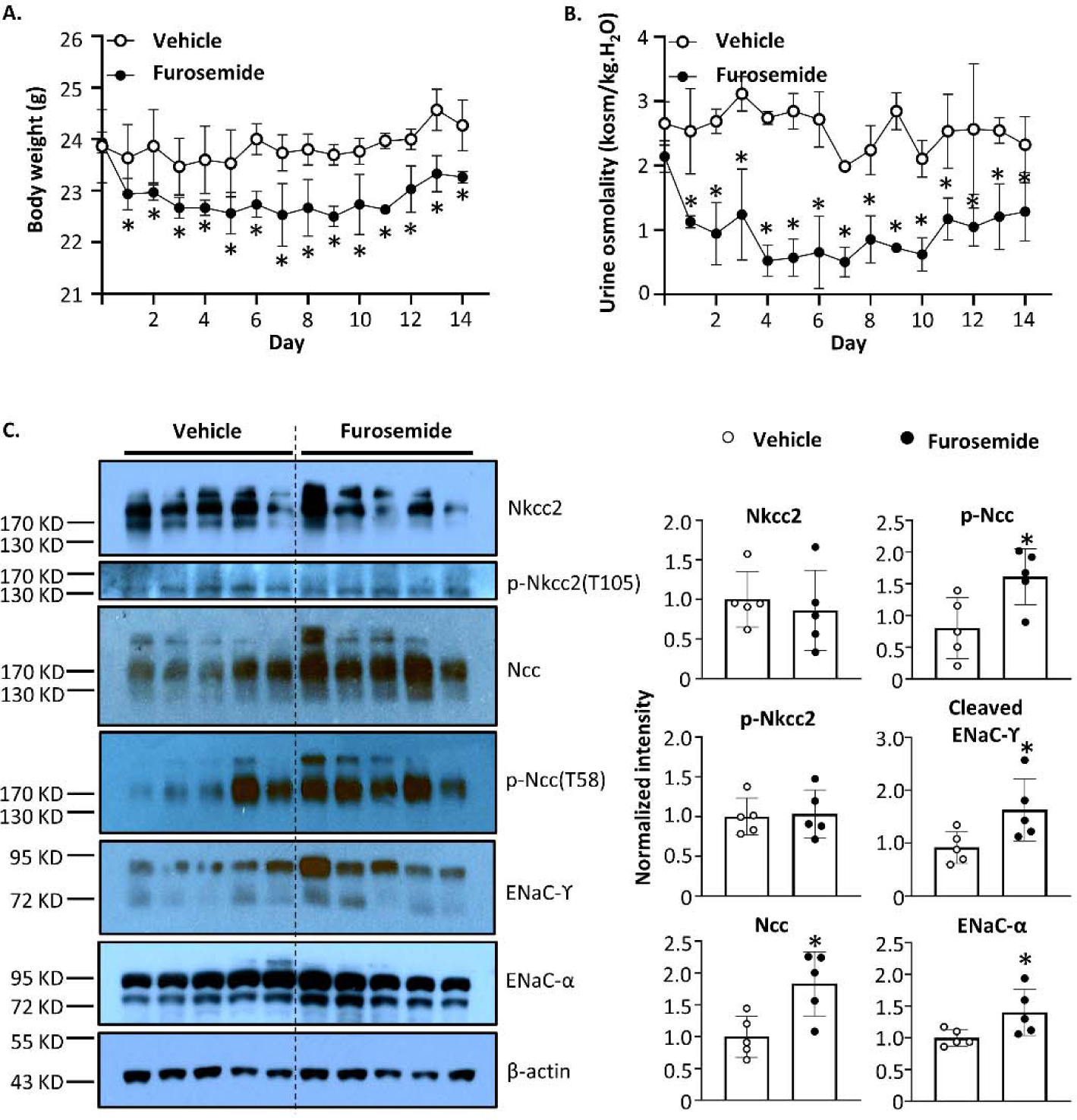
Characterization of 2-week furosemide-treated mice. **(A)** Daily body weight and **(B)** urine osmolality during 2-week furosemide or vehicle (DMSO) treatment. *p < 0.05 between vehicle-treated and furosemide-treated groups using two-way ANOVA with mixed-effects analysis (p values: body weight = 0.02, urine osmolality < 0.0001, n = 6 for each group). **(C)** Western blot analysis of major cotransporters and channels involved in sodium reabsorption in thick ascending limb, distal convoluted tubule, and collecting duct. The abundance of each band was measured by densitometry using the Image J program. *p < 0.05 between vehicle-treated and furosemide-treated groups using a two-tailed Student unpaired-t test (p values: Nkcc2 = 0.63, p-Nkcc2 = 0.84, Ncc = 0.01, p-Ncc = 0.02, cleaved ENaC-L = 0.04, ENaC-α = 0.05, n = 5 for each group). Data are presented as mean ± SD. (Ncc: sodium, chloride cotransporter; p-Ncc(T58): threonine 58 phosphorylated Ncc; Nkcc2: sodium, potassium, chloride cotransporter type 2; p-Nkcc2(T105): threonine 105 phosphorylated Nkcc2; ENaC: epithelial sodium channel)

TALs isolated from 2-week furosemide-treated mice had dramatically decreased basal (32.6±18.6 vs. 111.8±56.7pmol/min/cm, p=0.001), ATP-linked (14.0±6.2 vs. 36.7±15.0pmol/min/cm, p=0.007), and maximal (66.2±32.8 vs. 160.3±64.2pmol/min/cm, p=0.001), but not spare (34.2±25.3 vs. 58.2±33.9pmol/min/cm, p=0.44) mitochondrial respiration compared to vehicle-treated TALs (**Fig. 8A**, n=9). Two-hour ouabain treatment was used to acutely inhibit the sodium pump and active transport in TALs. Ouabain treatment lowered the basal (28.7±19.5 vs. 55.7±32.3pmol/min/cm, p=0.05) and ATP-linked (10.5±2.1 vs. 22.8±9.5pmol/min/cm, p=0.002) mitochondrial respiration in isolated TALs without changing maximal or spare mitochondrial respiratory capacity (**Fig. 8B**, n=9).

**Figure 8.**
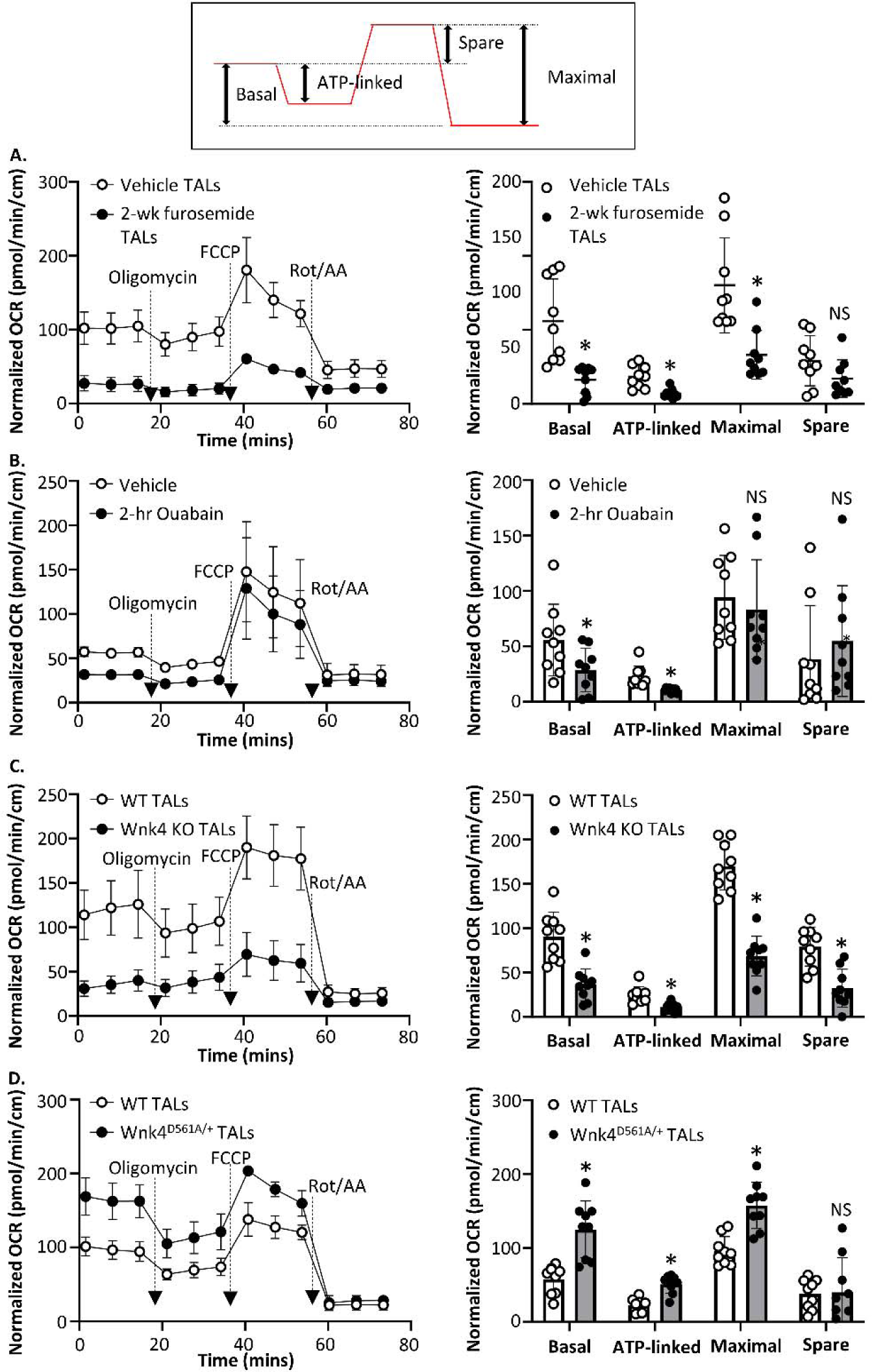
Mitochondrial respiration in isolated TALs with different transport activities. Seahorse XF Cell Mito Stress Tests were used to study mitochondrial respiration in isolated TALs with acutely or chronically altered transport activity. The inlet showed the definition of mitochondrial respiration parameters: Basal OCR = OCR_basal_ – OCR_Rot/AA_., ATP-linked = OCR_basal_ – OCR_Oligomycin_, Maximal OCR = OCR_FCCP_ – OCR_Rot/AA_, Spare OCR = OCR_FCCP_ – OCR_basal_. For each test, three TAL samples isolated from an experimental mouse were compared to three TAL samples isolated from a control mouse. Experimental and control mice were matched in genetic background, age, and gender. The left panel showed the OCR tracing of a representative test of three similar tests (see supplemental Fig. 4 for the other two tests). The dashed arrows indicate the injections of ETC complex inhibitors and uncoupler FCCP. All OCRs were normalized by tubule length. The right panel summarizes the results of mitochondrial respiration parameters from three tests. \**p* < 0.05, and NS: statistically not significant between two groups using a two-tailed Student unpaired-t test. Data are presented as mean ± SD. (**A**) TALs treated with 30mg/kg/day furosemide (2-wk furosemide) or vehicle for 2 weeks were compared (p values: basal = 0.001, ATP-linked = 0.007, maximal = 0.001, spare = 0.44, n=9). (**B**) TALs treated with 500 µM ouabain (2-hr Ouabain) or vehicle for 2 hours were compared. Tubule samples for each test were isolated from the same mouse kidney (p values: basal = 0.05, ATP-linked = 0.002, maximal = 0.59, spare = 0.49, n=9). (**C**) TALs isolated from Wnk4 knockout (Wnk4 KO) or wild-type (WT) littermates were compared (p values: basal = 0.0001, ATP-linked = 0.001, maximal = < 0.0001, spare = 0.003, n=9). (**D**) TALs isolated from Wnk4^D561A/+^ knockin or WT littermates were compared (p values: basal = 0.0002, ATP-linked = <0.0001, maximal = 0.0001, spare = 0.92, n=9).

Wnk4 kinase is known to enhance sodium reabsorption via sodium, chloride cotransporter (Ncc) and sodium, potassium, and chloride cotransporter type 2 (Nkcc2). Wnk4 knockout mice had a lower response to furosemide and thiazide, suggesting reduced endogenous Nkcc2 and Ncc activities (Terker et al., 2018). Conversely, gain-of-function D561A/+ Wnk4 mutation (mimicking human pseudohypoaldosteronism type II WNK4 mutation) enhanced the phosphorylation and activity of Nkcc2 and Ncc in vivo (Chu et al., 2013). Next, we studied the mitochondrial respiration in Wnk4 knockout and Wnk4^D561A/+^ knockin TALs/DCTs versus wild-type controls.

Wnk4 knockout TALs had lower basal (36.3±18.0 vs. 90.5±27.5pmol/min/cm, p=0.0001), ATP-linked (11.1±4.8 vs. 24.7±9.3pmol/min/cm, p=0.001), maximal (68.6±22.3 vs. 169.8±26.9pmol/min/cm, p<0.0001), and spare (32.4±21.5 vs. 79.3±22.0pmol/min/cm, p=0.0003) mitochondrial respiration than wild-type TALs (**Fig. 8C**, n=9). In contrast, Wnk4^D561A/+^ knockin TALs exhibited higher basal (125.2±38.5 vs. 57.7±19.1pmol/min/cm, p=0.0002), ATP-linked (50.9±12.0 vs. 22.7±9.0pmol/min/cm, p=<0.0001), and maximal mitochondrial respiration (157.6±31.3 vs. 95.9±19.5pmol/min/cm, p=0.0001) compared to controls (**Fig. 8D**, n=9). In DCTs, Wnk4 deletion caused a suppressed basal (21.1±14.2 vs. 73.9±20.8pmol/min/cm, p<0.0001), ATP-linked (10.0±6.8 vs. 27.0±8.5pmol/min/cm, p=0.0002), maximal (104.0±28.7 vs. 182.7±33.3pmol/min/cm, p<0.0001), and spare (80.2±29.2 vs. 112.1±33.9pmol/min/cm, p=0.048) mitochondrial respiration (**Fig. 9A**, n=9). Conversely, mitochondrial respiration was dramatically enhanced in Wnk4^D561A/+^ DCTs, including basal (127.6±24.4 vs. 64.6±12.8pmol/min/cm, p<0.0001), ATP-linked (48.2±13.6 vs. 28.4±10.9pmol/min/cm, p=0.004), and maximal capacity (191.8±49.4 vs. 100.1±16.2pmol/min/cm, p<0.0001) (**Fig. 9B**, n=9). Although 2-week furosemide treatment enhanced Ncc activities, the isolated DCTs from 2-week furosemide-treated mice showed only increased ATP-linked mitochondrial respiration (32.6±6.0 vs. 22.3±8.5pmol/min/cm, p=0.009) (**Fig. 9C**, n=9).

**Figure 9.**
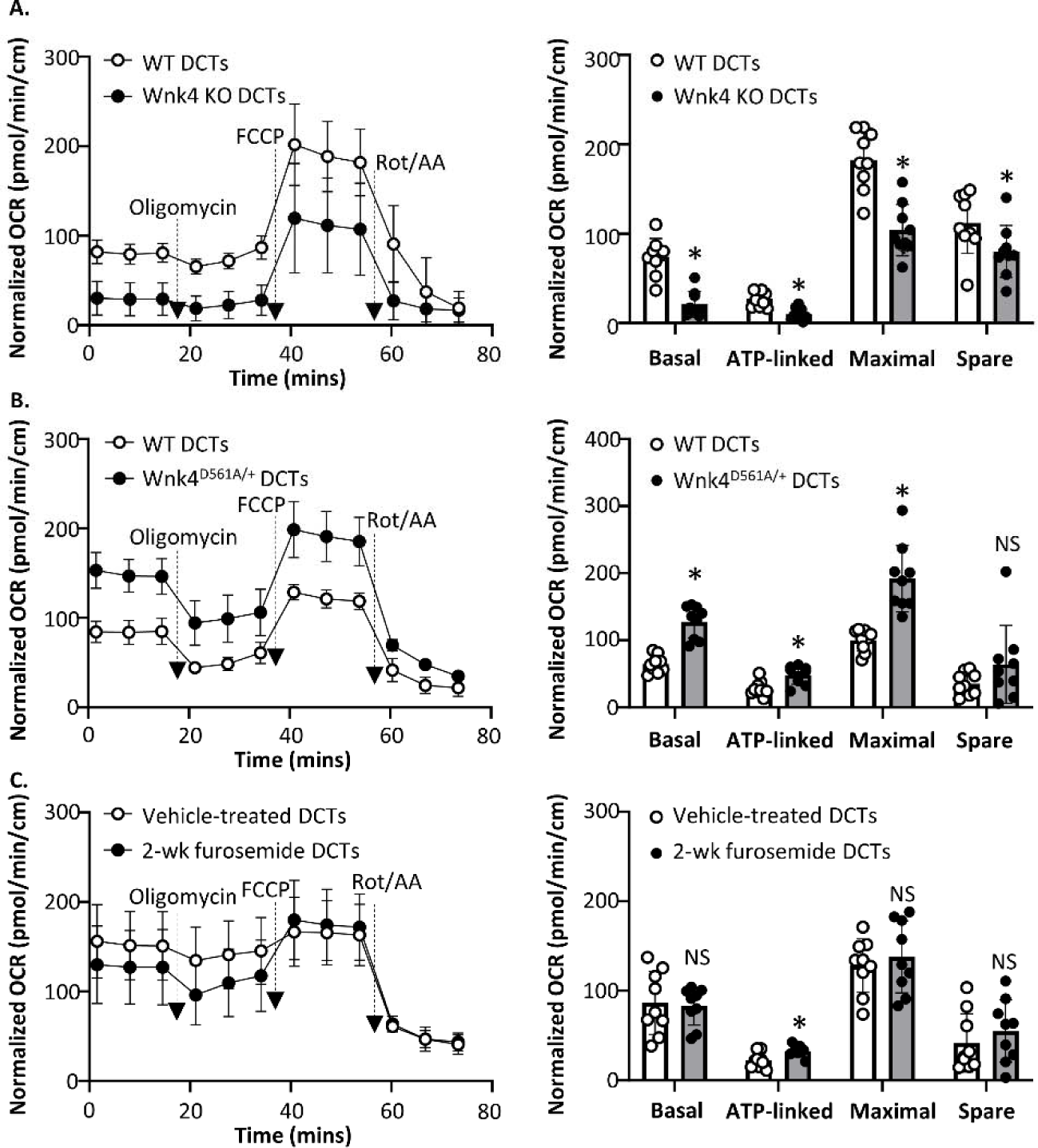
Mitochondrial respiration in isolated DCTs with different transport activities. Seahorse XF Cell Mito Stress Tests were used to study the mitochondrial respiration in isolated DCTs with chronically altered transport activity. For each test, three DCT samples isolated from an experimental mouse were compared to three TAL samples isolated from a control mouse. Experimental and control mice were matched in genetic background, age, and gender. The left panel showed the OCR tracing of a representative test of three similar tests (see supplemental Fig. 5 for the other two tests). The dashed arrows indicate the injections of ETC complex inhibitors and uncoupler FCCP. All OCRs were normalized by tubule length. The right panel summarizes the results of mitochondrial respiration parameters from three tests. \**p* < 0.05, and NS: statistically not significant between two groups using a two-tailed Student unpaired-t test. Data are presented as mean ± SD. (**A**) DCTs isolated from Wnk4 knockout (Wnk4 KO) or wild-type (WT) littermates were compared (p values: basal <0.0001, ATP-linked =0.0002, maximal <0.0001, spare = 0.05, n=9). (**B**) DCTs isolated from Wnk4^D561A/+^ knockin or WT controls were compared (p values: basal <0.0001, ATP-linked = 0.004, maximal <0.0001, spare = 0.17, n=9). (**C**) DCTs isolated from mice treated with 30mg/kg/day furosemide (2-wk furosemide) or vehicle were compared (p values: basal = 0.80, ATP-linked = 0.009, maximal = 0.58, spare = 0.42, n=9).

### Transport activity regulates mitochondrial mass and ultrastructure in TALs and DCTs

Next, we compared the mitochondrial morphology of isolated TALs and DCTs with reduced (Wnk4 knockout, furosemide-treated) or enhanced transport activity (Wnk4^D561A/+^) versus controls. The mitochondria in Wnk4 knockout TAL and DCT cells are fragmented and shorter (TALs: 1.13±0.60 vs. 1.66±0.65μm, p<0.0001; DCTs: 1.25±0.65 vs. 2.01±1.08μm, p<0.0001; n=50) and have lower mitochondrial volume density (TALs: 29.9±3.9 vs. 40.2±2.1%, p=0.0002; DCTs: 30.2±4.2 vs. 41.2±2.4%, p=0.0003; n=6) and cristae densities (the ratio of inner mitochondrial membrane to outer mitochondrial membrane, IMM/OMM)(TALs: 1.23±0.23 vs. 1.45±0.36, p=0.0009; DCTs: 1.23±0.33 vs. 1.45±0.38, p=0.002; n=50) than wild-type counterparts (**Fig. 10A, B**). Similarly, 2-week furosemide treatment decreases the mitochondrial volume density (35.9±4.4 vs. 49.4±8.5%, p=0.0003; n=10) and cristae density (IMM/OMM: 0.73±0.37 vs. 1.19±0.59, p<0.0001; n≥50) of TALs (**Fig. 10C**). In contrast, the mitochondria in Wnk4^D561A/+^ tubules tend to be longer (TALs: 1.88±0.79 vs. 1.66±0.65μm, p=0.13; DCTs: 3.08±1.38 vs. 2.01±1.08μm, p<0.0001; n=50) and have higher mitochondrial volume density (TALs: 46.1±2.4 vs. 40.2±2.1%, p=0.001; DCT: 50.7±1.8 vs. 41.2±2.4%, p<0.0001; n=6) than wild-type controls. The mitochondrial DNA (*Mt-nd1*) to nuclear DNA (*Hk1*) ratio was consistently decreased in loss-of-function groups (Wnk4 knockout TALs: 0.67±0.03 vs. 1.0±0.10, p=0.0008, n=4; Wnk4 knockout DCTs: 0.63±0.08 vs. 0.91±0.17, p=0.05, n=3; 2-week furosemide-treated TALs: 0.66±0.25 vs. 1.0±0.24, p=0.04, n=6) and statistically insignificant in gain-of-function groups (**Fig. 10D**).

**Figure 10.**
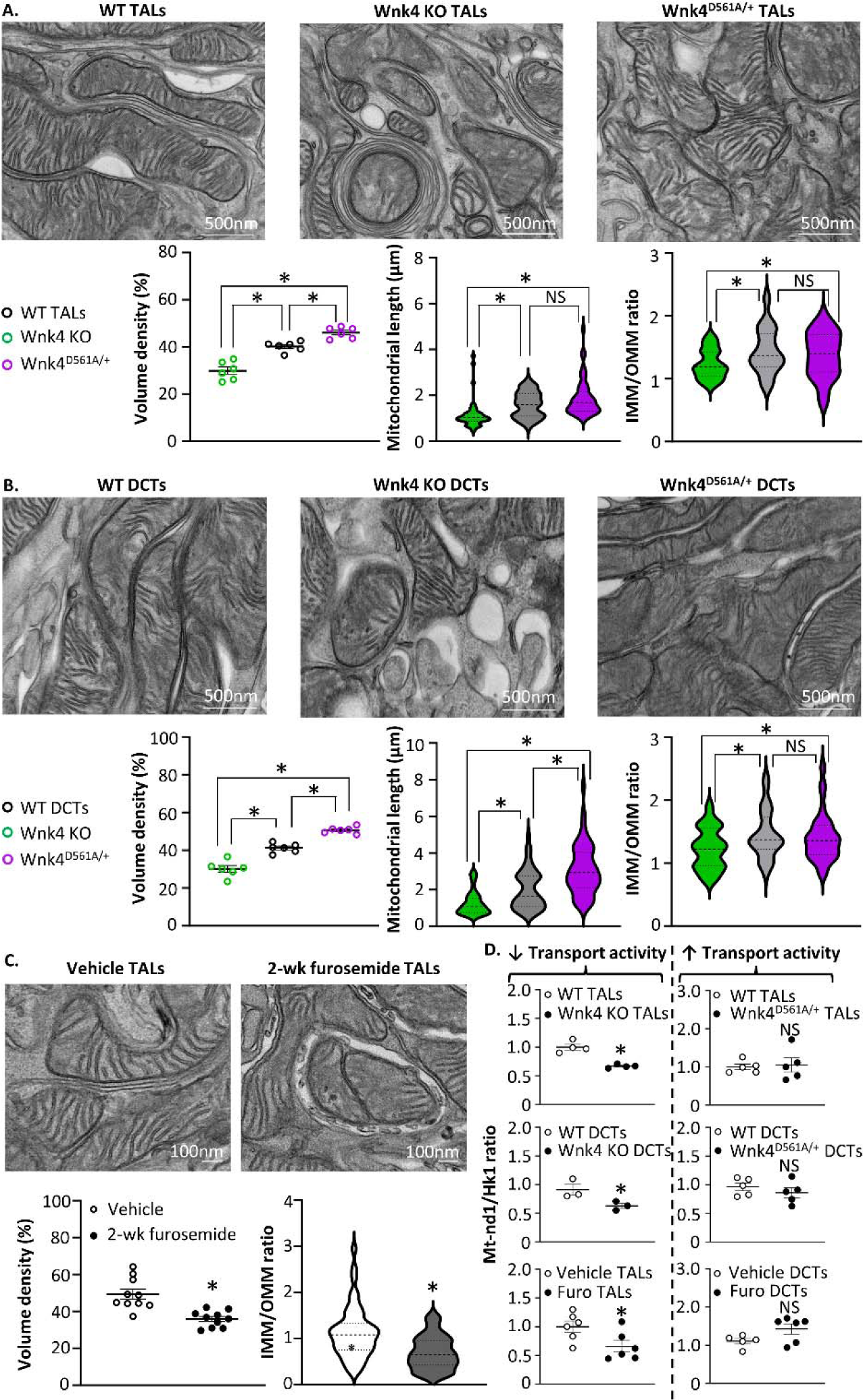
Mitochondrial morphology and DNA copy number of isolated TALs and DCTs with different transport activities. (**A, B**) Representative TEM images of TALs (**A**) and DCTs (**B**) isolated from Wnk4 knockout (Wnk4 KO), wild-type (WT), and Wnk4^D561A/+^ knockin (Wnk4^D561A/+^) mice. (**C**) Representative TEM images of isolated TALs from vehicle-treated (Vehicle) and 2-week furosemide-treated (2-wk furosemide) mice. Mitochondrial volume density (p values: TALs: Wnk4 KO vs. WT=0.0002; Wnk4^D561A/+^ vs. WT=0.0011; Wnk4 KO vs. Wnk4^D561A/+^<0.0001; DCTs: Wnk4 KO vs. WT =0.0003; Wnk4^D561A/+^ vs. WT<0.0001; Wnk4 KO vs. Wnk4^D561A/+^<0.0001; Vehicle vs. 2-wk furosemide: 0.003; n=6 tubules for A, B; n=10 for C), length (p values: TALs: Wnk4 KO vs. WT <0.0001; Wnk4^D561A/+^ vs. WT =0.13; Wnk4 KO vs. Wnk4^D561A/+^<0.0001; DCTs: Wnk4 KO vs. WT<0.0001; Wnk4^D561A/+^ vs. WT<0.0001; Wnk4 KO vs. Wnk4^D561A/+^<0.0001; n=50 mitochondria), and cristae density (p values: TALs: Wnk4 KO vs. WT=0.0009; Wnk4^D561A/+^ vs. WT=0.36; Wnk4 KO vs. Wnk4^D561A/+^=0.02; DCTs: Wnk4 KO vs. WT=0.002; Wnk4^D561A/+^ vs. WT=0.59; Wnk4 KO vs. Wnk4^D561A/+^=0.01; Vehicle vs. 2-wk furosemide<0.0001; n=50 mitochondria) were analyzed using Image J (see methods). (see the entire raw data in Supplemental Table 1-3) (**D**) The mitochondrial DNA (*Nd1* gene) to nuclear DNA (*Hk1* gene) ratios in isolated TALs or DCTs with decreased or increased transport activity. \**p* < 0.05, and NS: statistically not significant between two groups using a two-tailed Student unpaired-t test (p values: WT TALs vs. Wnk4 KO TALs=0.0008; WT DCTs vs. Wnk4 KO DCTs=0.05, Vehicle TALs vs. Furo TALs=0.04, WT TALs vs. Wnk4^D561A/+^ TALs=0.80, WT DCTs vs. Wnk4^D561A/+^ DCTs=0.38, Vehicle DCTs vs. Furo DCTs=0.08; n≥3). Data are presented as mean ± SD. (Furo: furosemide; IMM: inner mitochondrial membrane; OMM: outer mitochondrial membrane; TEM: transmission electron microscopy)

## Discussion

This study demonstrates that the EFA assay can be applied to investigate mitochondrial respiration and energy metabolism in freshly micro-dissected renal tubules. Coupled with genetic or pharmacological manipulations, the assay is a powerful tool capable of providing segment-specific mitochondrial metabolism. This approach is superior to assays using bulk kidney tissues or culture cells by preserving the characters and microenvironment of renal tubular cells and allowing real-time measurements of oxygen consumption and acid production.

One major advantage of our experiment system and assay is robust metabolic activity. In the assay, each tubular sample contains an estimated only hundreds of cells, an order of magnitude lower in number than required when renal tubular cells are grown in culture (≥5000 cells in culture generally recommended). This is because renal tubular cells are rich in mitochondria, and, more importantly, cells are metabolically active when preserved in freshly isolated renal tubules. Interestingly, we found that increasing tubule size beyond 1.5-2 cm in length may exceed oxygen supply in the assay system, leading to increased anaerobic glycolysis over mitochondrial OXPHOS. In addition, high basal mitochondrial respiration may compromise the instrument’s ability to measure the maximal and spare mitochondrial capacity due to reduced responsivity to the uncoupler FCCP. Therefore, the optimal starting sample size is < 1.4 cm for TALs/DCTs and < 1.0 cm for PTs and can be adjusted to the basal OCRs within the range of 50-150 pmol/min and ECARs ∼10 mpH/min.

Having established and validated the assay, we further determine the metabolic characteristics of PTs, TALs, and DCTs. Isolated PTs are highly oxidative and relatively deficient in glycolysis, supported by the lack of compensatory ECARs elevation when ETC complex inhibitors shut off OXPHOS. These findings are consistent with previous reports of low glycolytic enzyme expression and activity and little lactate formation in PTs (Dickman & Mandel, 1989; Guder & Ross, 1984; Guder et al., 1986; Uchida & Endou, 1988). In contrast, TALs and DCTs have high endogenous glycolytic capability and significant anaerobic glycolysis. We further examined substrate utilization. Previous studies suggest that TALs and DCTs can metabolize a slew of substrates with unclear relative importance to ATP production (Klein et al., 1981; Wittner et al., 1984; Chamberlin & Mandel, 1986; Wirthensohn & Guder, 1986; Uchida & Endou, 1988). By using metabolic pathway inhibitors, we delineate the physiological importance of these energy pathways. Our results provide compelling functional evidence that glucose is the primary energy source for TALs and DCTs in basal and high-energy demand states. Previous studies have shown that glutaminase is highly expressed in TALs and DCTs (Guder & Ross, 1984), and TALs generated ^14^C-labeled CO_2_ when provided with isotope-labeled glutamine in vitro (Klein et al., 1981), implying that TALs and DCTs may utilize glutamine as an energy substrate. However, our results showed that isolated TALs and DCTs had a relatively small response to glutaminase inhibitor BPTES compared to MPC inhibitor UK-5099, indicating that glutaminolysis is a minor pathway for OXPHOS in these tubules. The enzymes for β-oxidation of fatty acids are present in all kidney regions, with the highest activities in PTs and DCTs (Le Hir M & Dubach UC, 1982). Our results demonstrate that LCFA oxidation fuels DCTs in the basal state but is not essential for the high oxygen consumption state. TALs do not oxidize LCFAs for ATP production when glucose is available. Our results yet do not rule out the importance of short-chain fatty acids (SCFAs) (e.g., butyrate, acetate) that may fuel salt reabsorption in TALs (Wittner et al., 1984). SCFAs enter mitochondria independently from CPT1; therefore, etomoxir cannot discern the contribution of SCFAs to OXPHOS.

Most ATP production (∼70%) in renal tubules is consumed by active transport (Harris et al., 1981; Mandel, 1986). Early studies identified a linear correlation between oxygen consumption and sodium reabsorption rates in perfused dog kidneys (Thurau 1961). Ouabain, a sodium pump inhibitor, reduced oxygen consumption significantly (50-70%) in renal cortical tubule suspension (Balaban et al., 1980a; 1980b). In humans, furosemide quickly increased medullary oxygenation, probably due to diverting salt reabsorption in TALs to downstream cortical segments (Prasad et al., 1996; Haddock et al., 2019). Mitochondrial gene mutations have been linked to Gitelman-like syndrome with impaired active transport in DCTs, probably by disrupting OXPHOS (Viering et al., 2022). Overall, the results support the notion of coupling between active transport and mitochondrial respiration in TALs and DCTs.

Whittam model has been used to explain how transport is coupled to mitochondrial metabolism (Whittam 1961; Mandel & Balaban, 1981): acute suppression of transport activity either by increasing intracellular ATP or ATP/ADP ratio feeds back to downregulate ATP production from mitochondria. In this study, we found that acute ouabain treatment reduced the basal OCR of TALs by 60%. Most of this reduction occurs at the level of basal and ATP-linked respiration revealed by oligomycin-sensitive respiration (before and after oligomycin that inhibits ATP synthase). The results support Whittam model that ATP production is reduced when transport activity and energy demand are lowered in renal tubules. The maximal mitochondrial capacity is not affected in the model of acute transport inhibition.

We further study mitochondrial adaptation to chronic alteration of transport activity in renal tubules. In addition to the mitochondrial respiration parameters, we measured mitochondrial morphologies and mitochondrial DNA copy numbers in the chronic models. Unlike acute suppression of transport activity using ouabain, tubules with chronic suppression of transport, including furosemide-treated TALs and Wnk4-deleted TALs/DCTs, consistently display remarkably decreased maximal mitochondrial capacity, apart from reduced basal and ATP-linked respirations. Furthermore, they exhibit similar mitochondrial morphological changes (e.g., reduced mitochondrial length, volume density, cristae density) and low mitochondrial DNA copy number, supporting the idea that chronic transport suppression downregulates mitochondrial biogenesis. Three gain-of-function models were studied, including furosemide-treated DCTs and Wnk4D561A/+ TALs/DCTs. Wnk4^D561A/+^ TALs and DCTs had higher basal respiration, ATP-linked respiration, and maximal respiratory capacity than controls. Morphologically, the TEM images of Wnk4^D561A/+^ TALs and DCTs showed increased mitochondrial length and volume density. However, the mitochondrial DNA copy numbers in gain-of-function groups are not significantly different from controls, indicating that non-transcriptional regulations, such as mitochondrial fusion, may be the underlying mechanisms (Bennett CF et al., 2022). Two-week furosemide treatment only enhanced the ATP turnover rate in DCTs without changing the maximal capacity. Therefore, the etiology and duration of high transport activity may determine how mitochondria adapt to long-term transport activity.

In conclusion, we have established a reliable new method to study mitochondrial function and metabolism in isolated renal tubules exhibiting high metabolic activity. Using the methodology, we have begun to dissect the mechanism of coupling between transport activity and renal tubular remodeling, known as “disuse atrophy” and “use hypertrophy”. The mechanisms underlying tubular remodeling adapting to transport activity alteration remain largely unknown. For example, changes in intracellular concentration of small molecules (e.g., ATP/ADP, ions), while they may be the signals for regulating ATP synthase turnover rate in the acute setting, are unlikely to be responsible for chronic mitochondrial adaptation because these acute changes should be normalized when ATP synthase or transepithelial transport is balanced. Furthermore, it is unknown whether and how alterations in mitochondrial function and metabolism mediate changes in cell proliferation. Future studies correlating single-tubule mitochondrial metabolism with multimodal omics may be needed to fill the knowledge gap, which may help unlock the molecular mechanisms of congenital tubulopathies and acquired renal tubule remodeling.

## Additional information

### Data availability statement

The data that support the findings of this study are available from the corresponding author upon reasonable request.

### Competing interests

The authors have no competing interests to disclose.

### Author Contributions

C.-J.C. designed research works, conducted experiments, acquired data, and analyzed data; D.-D.F., N.-J.M., and C.-L.H. revised the manuscript critically for important intellectual content; C.-J.C. drafted and finalized the paper; all authors approved the final version of the manuscript, agree to be accountable for all aspects of the work in ensuring that questions related to the accuracy or integrity of any part of the work are appropriately investigated and resolved.

### Funding

This study is supported by grants from the National Institutes of Health, USA (DK134420 to CJC; DK111542 to CLH).

## Supporting information

Additional supporting information can be found online in the Supporting Information section at the end of the HTML view of the article.

## Abbreviations

BPTES: glutaminase inhibitor Bis-2-(5-phenylacetamido-1,3,4-thiadiazol-2-yl)ethyl sulfide
CPT1: carnitine palmitoyltransferase I
DCT: distal convoluted tubule
ECAR: extracellular acidification rate
EFA: extracellular flux analysis
ETC: electron transport chain
FCCP: mitochondrial uncoupler carbonyl cyanide-p-trifluoromethoxyphenylhydrazone
Hk1: hexokinase 1
LCFA: long-chain fatty acid
Mt-nd1: mitochondrial NADH-ubiquinone oxidoreductase chain 1
Ncc: sodium, chloride cotransporter
Nkcc2: sodium, potassium, and chloride cotransporter type 2
OCR: oxygen consumption rate
OXPHOS: oxidative phosphorylation
PER: proton efflux rate
PT: proximal tubule
Rot/AA: ETC complex inhibitor rotenone/antimycin A
SCFA: short-chain fatty acid
TAL: thick ascending limb
TCA: tricarboxylic acid
TEM: transmission electron microscope
Wnk4: with-no-lysine kinase 4

## Notes

### Competing Interest Statement

The authors have declared no competing interest.

