## Supplemental materials for "Transport activity regulates mitochondrial bioenergetics and biogenesis in renal tubules"

**Supporting information**

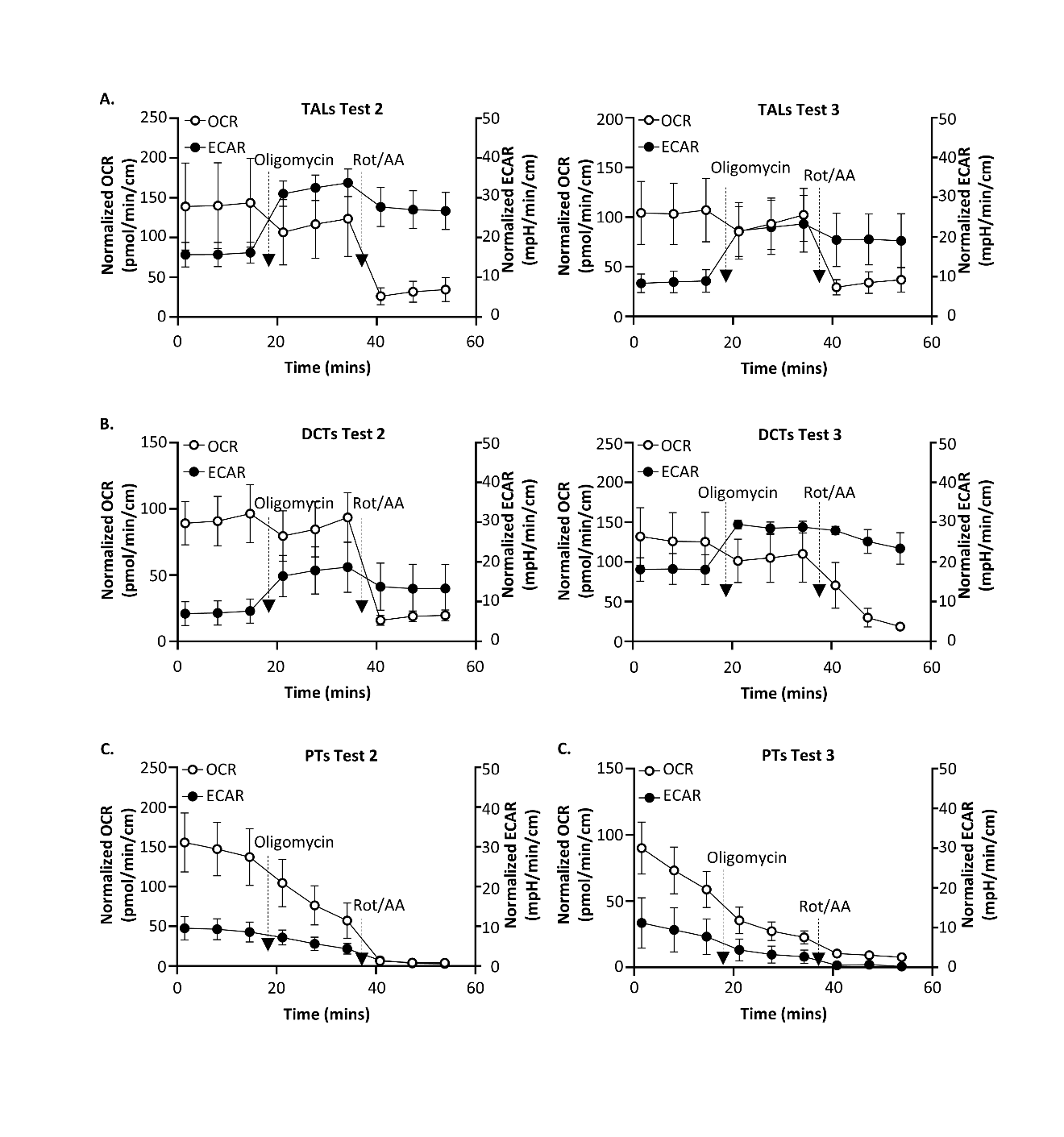

**Supplemental Figure 1. ATP production rate assays for isolated renal tubules.** Additional tests for Fig. 4A**.** Real-time ATP rate tests were used to determine the relative contributions of mitochondrial OXPHOS and glycolysis to ATP production. Simultaneous OCRs and ECARs measurements in real-time ATP rate tests using isolated TALs (**A**), DCTs (**B**), or PTs (**C**)

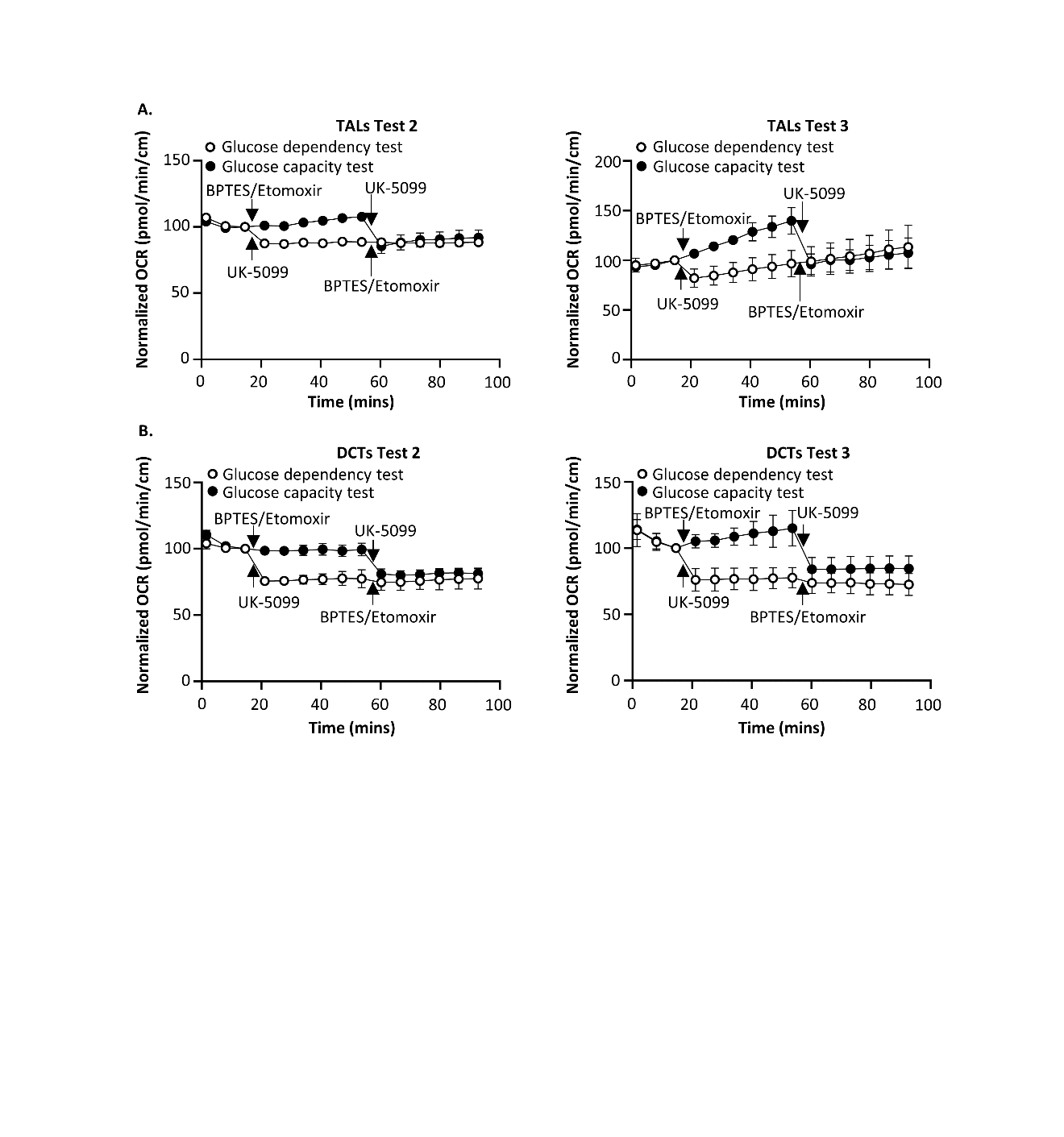

**Supplemental Figure 2. Mito Fuel Flex tests for TALs and DCTs.** Additional tests for Fig. 5A, C. Seahorse XF Mito Fuel Flex Tests were used to determine the relative contributions of glucose, glutamine, and LCFAs oxidation to basal respiration in isolated TALs (**A**) and DCTs (**B**).

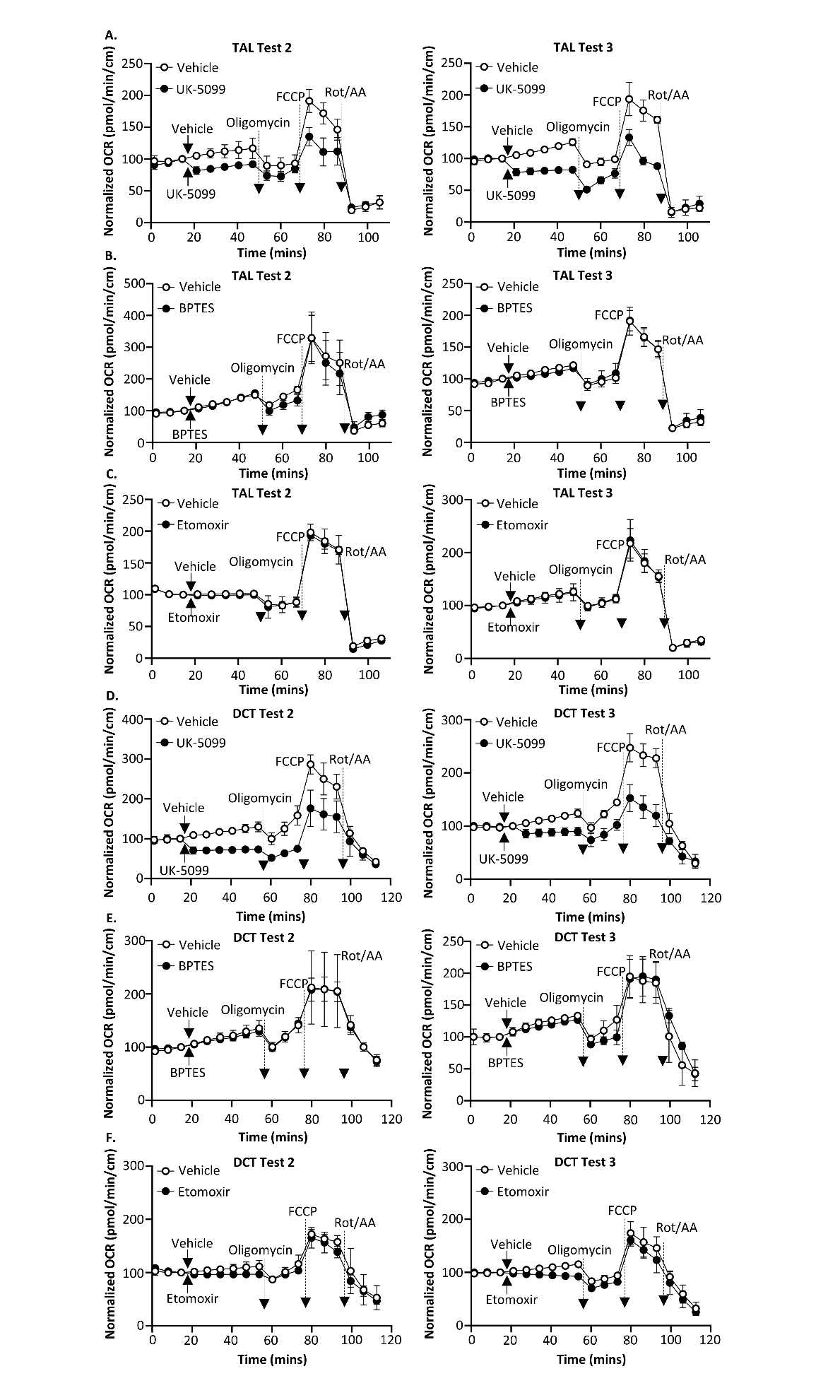
**Supplemental Figure 3. Substrate oxidation stress tests for TALs and DCTs.** Additional tests for Fig. 6. Seahorse XF Substrate Oxidation Stress Tests were conducted to investigate the oxidation of glucose, glutamine, and LCFAs in basal and high energy demand states in TALs (**A-C**) and DCTs (**D-F**).

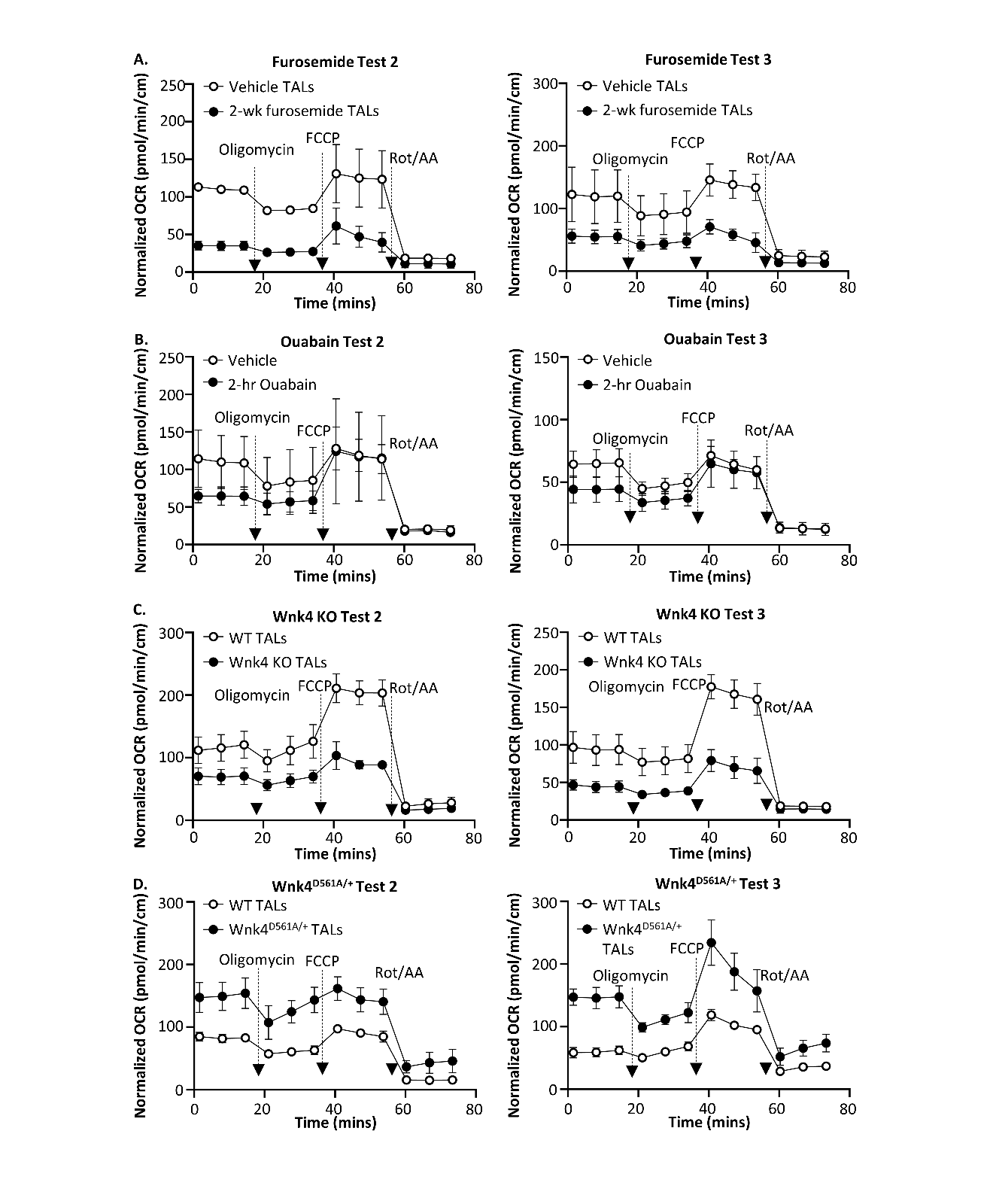
**Supplemental Figure 4. Mitochondrial respiration in isolated TALs with different transport activities.** Additional tests for Fig. 8. Seahorse XF Cell Mito Stress Tests were used to study mitochondrial respiration in isolated TALs with acutely or chronically altered transport activity.

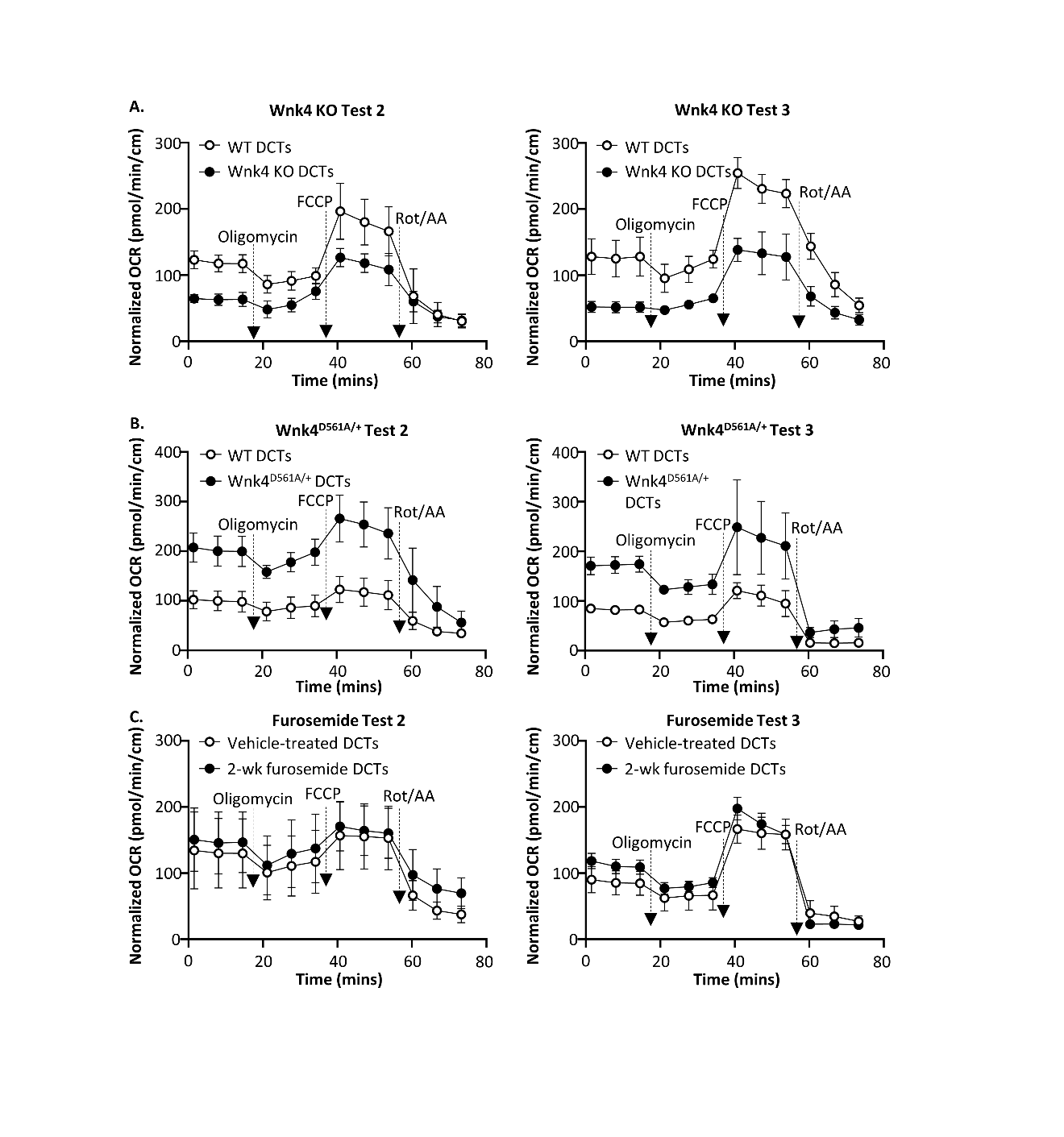
**Supplemental Figure 5. Mitochondrial respiration in isolated DCTs with different transport activities.** Additional tests for Fig. 9. Seahorse XF Cell Mito Stress Tests were used to study the mitochondrial respiration in isolated DCTs with chronically altered transport activity.

**Supplemental Table 1.** **Mitochondrial morphology of isolated TALs with different transport activities.** The entire raw dataset of Fig. 10A.

| **Mitochondrial length (μm)** | | | **IMM/OMM ratio** | | |
| --- | --- | --- | --- | --- | --- |
| **WNK4 KO**  **(n=50)** | **WT**  **(n=50)** | **WNK4^D561A/+^**  **(n=50)** | **WNK4 KO**  **(n=50)** | **WT**  **(n=50)** | **WNK4^D561A/+^**  **(n=50)** |
| 2.09 | 1.06 | 3.82 | 1.42 | 1.35 | 1.9 |
| 0.97 | 1.05 | 1.31 | 1.17 | 1.52 | 1.52 |
| 1.2 | 0.91 | 2.1 | 0.99 | 0.87 | 0.85 |
| 2.38 | 1.45 | 2.92 | 1.22 | 1.21 | 1.78 |
| 2.04 | 1.03 | 1.41 | 1.7 | 1.25 | 1.71 |
| 0.75 | 0.95 | 1.42 | 1.46 | 1.77 | 1.12 |
| 0.59 | 1.04 | 1.7 | 1.13 | 0.92 | 1.92 |
| 1.36 | 0.61 | 1.52 | 1.12 | 1.36 | 0.72 |
| 2.48 | 0.88 | 1.74 | 1.05 | 1.08 | 1.76 |
| 2.46 | 0.68 | 3.18 | 1.28 | 1.23 | 0.78 |
| 0.95 | 0.96 | 1.38 | 1.2 | 1.11 | 0.57 |
| 1.59 | 0.68 | 1.97 | 1.42 | 1.31 | 0.99 |
| 1.07 | 0.55 | 1.51 | 1.35 | 1.35 | 1.57 |
| 1.68 | 1.19 | 1.06 | 1.3 | 2.27 | 1.59 |
| 1.11 | 2.55 | 1.18 | 1.25 | 1.39 | 1.4 |
| 0.86 | 1.55 | 1.84 | 1.14 | 2.21 | 1.51 |
| 1.69 | 1.04 | 4.89 | 1.05 | 1.59 | 1.62 |
| 2.28 | 3.71 | 1.84 | 1.18 | 1.18 | 1.83 |
| 2.1 | 3.38 | 2.53 | 1.2 | 1.46 | 1.62 |
| 1.56 | 0.81 | 1.75 | 1.46 | 1.26 | 2.07 |
| 2.71 | 0.46 | 1.34 | 1.13 | 1.83 | 1.3 |
| 1.85 | 1.09 | 2.78 | 1.41 | 2 | 1.95 |
| 1.12 | 0.86 | 3.79 | 0.93 | 0.92 | 1.31 |
| 0.89 | 1.21 | 1.94 | 1.6 | 1.89 | 1.7 |
| 1.11 | 0.91 | 1.74 | 1.16 | 1.15 | 1.06 |
| 1.93 | 1.15 | 1.59 | 0.9 | 0.82 | 1.23 |
| 1.14 | 1.26 | 2.35 | 1.13 | 1.48 | 0.94 |
| 0.82 | 0.94 | 1.26 | 0.98 | 1.3 | 0.98 |
| 1.62 | 1.15 | 1.5 | 0.92 | 1.27 | 1.14 |
| 1.82 | 0.65 | 1.71 | 1.59 | 1.19 | 1.71 |
| 1.15 | 1.61 | 1.32 | 0.98 | 1.68 | 1.26 |
| 2.42 | 1.25 | 1.92 | 1.56 | 1.54 | 1.13 |
| 2.17 | 0.67 | 1.04 | 1.07 | 1.01 | 1.16 |
| 2.1 | 0.94 | 1.58 | 1.21 | 1.91 | 1.79 |
| 2.5 | 0.7 | 1.32 | 1.68 | 1.72 | 0.97 |
| 2.05 | 1.27 | 1.21 | 1.42 | 1.38 | 1.12 |
| 1.33 | 0.7 | 1.29 | 0.93 | 1.37 | 1.55 |
| 1.2 | 0.95 | 2.44 | 1.03 | 1.04 | 1.03 |
| 1.16 | 1.14 | 1.45 | 0.97 | 1.67 | 1.75 |
| 0.87 | 1.21 | 1.68 | 1.46 | 1.51 | 1.87 |
| 2.34 | 1.06 | 2.11 | 1.05 | 1.79 | 1.47 |
| 0.95 | 1.37 | 1.05 | 1.53 | 1.66 | 1.7 |
| 1.22 | 1.02 | 1.23 | 1.23 | 1.3 | 1.5 |
| 1.81 | 0.78 | 1.3 | 1.58 | 0.97 | 1.4 |
| 1.56 | 1.04 | 2.29 | 1.03 | 1.38 | 1.28 |
| 1.88 | 0.92 | 2.32 | 1.65 | 1.81 | 1.09 |
| 2.08 | 0.95 | 3 | 1.11 | 2.34 | 1.38 |
| 1.81 | 0.92 | 1.94 | 0.85 | 1.18 | 1.27 |
| 1.59 | 0.97 | 1.38 | 1.02 | 1.86 | 0.64 |
| 1.56 | 1.42 | 1.14 | 1.42 | 1.24 | 1.39 |

**Supplemental Table 2.** **Mitochondrial morphology of isolated DCTs with different transport activities.** The entire raw dataset of Fig. 10B.

| **Mitochondrial length (μm)** | | | **IMM/OMM ratio** | | |
| --- | --- | --- | --- | --- | --- |
| **WNK4 KO**  **(n=50)** | **WT**  **(n=50)** | **WNK4^D561A/+^**  **(n=50)** | **WNK4 KO**  **(n=50)** | **WT**  **(n=50)** | **WNK4^D561A/+^**  **(n=50)** |
| 0.87 | 1.66 | 4.30 | 1.19 | 2.37 | 1.07 |
| 0.66 | 1.45 | 3.88 | 1.71 | 1.08 | 1.62 |
| 0.68 | 0.85 | 5.53 | 1.39 | 1.45 | 2.47 |
| 0.63 | 0.64 | 3.16 | 1.77 | 2.25 | 2.08 |
| 1.31 | 2.44 | 3.22 | 1.00 | 1.01 | 1.40 |
| 1.08 | 0.75 | 2.47 | 0.93 | 1.10 | 1.23 |
| 0.78 | 2.80 | 4.45 | 1.55 | 1.88 | 1.05 |
| 0.97 | 2.00 | 4.35 | 0.83 | 0.80 | 0.81 |
| 1.41 | 2.60 | 4.68 | 0.95 | 1.00 | 1.27 |
| 0.67 | 2.10 | 2.98 | 1.02 | 1.41 | 1.35 |
| 1.74 | 3.06 | 6.16 | 1.14 | 1.34 | 1.21 |
| 0.76 | 1.08 | 4.27 | 1.24 | 1.41 | 1.38 |
| 1.53 | 3.49 | 4.31 | 1.60 | 1.02 | 1.32 |
| 0.76 | 1.28 | 2.12 | 1.60 | 1.89 | 0.86 |
| 0.73 | 0.87 | 2.15 | 1.00 | 1.00 | 1.60 |
| 2.56 | 1.31 | 3.10 | 1.31 | 0.90 | 1.47 |
| 1.07 | 1.53 | 1.33 | 0.72 | 1.34 | 1.37 |
| 1.95 | 1.02 | 2.68 | 1.60 | 1.38 | 1.33 |
| 0.67 | 1.61 | 2.49 | 0.89 | 2.20 | 1.03 |
| 0.65 | 1.72 | 2.96 | 1.64 | 0.93 | 1.08 |
| 1.20 | 2.82 | 2.16 | 0.82 | 1.25 | 1.04 |
| 1.19 | 3.02 | 2.35 | 1.31 | 1.32 | 1.49 |
| 1.20 | 1.20 | 3.81 | 1.34 | 1.39 | 1.56 |
| 0.93 | 0.78 | 3.97 | 1.56 | 1.37 | 1.41 |
| 0.92 | 1.03 | 2.62 | 1.30 | 1.30 | 1.39 |
| 1.48 | 1.36 | 1.25 | 0.87 | 1.23 | 1.10 |
| 0.85 | 1.62 | 3.50 | 0.82 | 1.42 | 1.65 |
| 1.07 | 0.89 | 1.41 | 1.63 | 1.54 | 2.57 |
| 1.71 | 1.21 | 3.53 | 0.96 | 1.00 | 2.24 |
| 1.34 | 2.75 | 2.62 | 1.59 | 1.62 | 0.86 |
| 1.08 | 1.07 | 2.47 | 0.79 | 1.81 | 1.69 |
| 2.62 | 2.71 | 2.03 | 0.71 | 1.20 | 1.17 |
| 2.93 | 2.83 | 3.50 | 1.04 | 1.38 | 0.96 |
| 0.75 | 0.93 | 1.59 | 0.96 | 1.18 | 2.22 |
| 1.19 | 2.64 | 2.57 | 1.90 | 1.72 | 1.41 |
| 0.86 | 2.66 | 2.91 | 1.36 | 1.99 | 1.31 |
| 0.70 | 0.68 | 1.47 | 1.35 | 2.36 | 1.89 |
| 1.63 | 4.25 | 1.69 | 1.14 | 1.36 | 1.46 |
| 0.50 | 1.48 | 2.00 | 0.95 | 1.97 | 1.19 |
| 0.64 | 1.18 | 2.11 | 1.17 | 1.25 | 1.43 |
| 1.64 | 0.79 | 3.00 | 1.25 | 1.93 | 1.14 |
| 0.58 | 3.31 | 7.65 | 0.95 | 1.52 | 1.62 |
| 2.23 | 1.42 | 5.00 | 1.25 | 1.33 | 1.08 |
| 0.91 | 2.64 | 4.34 | 1.92 | 1.30 | 1.25 |
| 0.87 | 3.26 | 4.79 | 1.20 | 1.79 | 1.20 |
| 1.52 | 4.95 | 2.55 | 1.29 | 1.60 | 1.47 |
| 1.10 | 3.82 | 3.17 | 0.72 | 1.28 | 1.77 |
| 2.89 | 4.24 | 1.23 | 1.59 | 1.29 | 1.19 |
| 1.84 | 2.05 | 1.05 | 1.40 | 1.40 | 1.12 |
| 2.83 | 2.63 | 1.24 | 1.20 | 1.80 | 1.64 |

**Supplemental Table 3.** **Mitochondrial morphology of isolated TALs from vehicle-treated (Vehicle) and 2-week furosemide-treated.** The entire raw dataset of Fig. 10C.

| **IMM/OMM ratio** | |
| --- | --- |
| **Vehicle** | **2-wk furosemide** |
| 1.73 | 0.27 |
| 1.00 | 0.62 |
| 0.92 | 0.77 |
| 0.98 | 0.78 |
| 0.49 | 0.25 |
| 1.94 | 0.50 |
| 1.20 | 0.66 |
| 1.35 | 0.24 |
| 0.79 | 0.23 |
| 1.58 | 0.79 |
| 1.18 | 0.46 |
| 2.51 | 0.85 |
| 1.74 | 0.65 |
| 0.56 | 0.45 |
| 0.71 | 0.45 |
| 0.85 | 0.72 |
| 2.31 | 1.33 |
| 1.18 | 1.10 |
| 1.50 | 0.57 |
| 1.32 | 0.28 |
| 1.56 | 0.38 |
| 2.95 | 0.17 |
| 1.20 | 0.21 |
| 0.63 | 0.55 |
| 1.32 | 0.44 |
| 1.98 | 0.65 |
| 0.78 | 0.93 |
| 0.75 | 1.14 |
| 1.11 | 0.34 |
| 0.36 | 0.99 |
| 1.10 | 1.05 |
| 0.88 | 1.50 |
| 1.22 | 1.43 |
| 0.30 | 0.23 |
| 1.32 | 0.66 |
| 1.12 | 0.44 |
| 1.56 | 0.37 |
| 0.48 | 0.49 |
| 1.03 | 0.93 |
| 0.60 | 1.27 |
| 1.25 | 1.47 |
| 0.98 | 0.62 |
| 0.92 | 0.93 |
| 1.13 | 0.15 |
| 0.46 | 1.04 |
| 0.69 | 0.78 |
| 0.27 | 0.96 |
| 0.85 | 0.56 |
| 0.75 | 0.95 |
| 1.07 | 1.10 |
